# A rapidly reversible mutation generates subclonal genetic diversity and unstable drug resistance

**DOI:** 10.1101/2020.03.03.972455

**Authors:** Lufeng Dan, Yuze Li, Shuhua Chen, Jingbo Liu, Fangting Li, Yu Wang, Xiangwei He, Lucas B. Carey

## Abstract

Most genetic changes have negligible reversion rates. As most mutations that confer resistance to an adversary condition (e.g., drug treatment) also confer a growth defect in its absence, it is challenging for cells to genetically adapt to transient environmental changes. Here we identify a set of rapidly reversible drug resistance mutations in *S. pombe* that are caused by Microhomology mediated Tandem Duplication (MTD), and reversion back to the wild-type sequence. Using 10,000x coverage whole-genome sequencing we identify near 6000 subclonal MTDs in a single clonal population, and determine using machine learning how MTD frequency is encoded in genome. We find that sequences with the highest predicted MTD rates tend to generate insertions that maintain the correct reading frame suggesting that MTD formation has shaped the evolution of coding sequences. Our study reveals a common mechanism of reversible genetic variation that is beneficial for adaptation to environmental fluctuations and facilitates evolutionary divergence.

## Main Text

Different mechanisms of adaptation have different timescales. Epigenetic changes are often rapid and reversible, while most genetic changes have nearly negligible rates of reversion(1). This poses a challenge for genetic adaptation to transient conditions such as drug treatment; mutations that confer drug resistance are often deleterious in the absence of drug, and the second-site suppressor mutations are required to restore fitness(2, 3). Pre-existing tandem repeats (satellite DNA) undergo frequent expansion and contraction (4–6). While repeats are rare inside of most coding sequences and functional elements, there is some evidence for conserved repetitive regions that undergo expansion and contraction to regulate protein functions or expression(6–8). RNAi- or Chromatin-based epigenetic states have been associated with transient drug resistance in fungi(9) and cancer cells(10, 11), and transiently resistant states have been characterized by differences in organelle state, growth rate, and gene expression in budding yeast(12, 13). In bacteria and in fungi, copy-number gain and subsequent loss can result in reversible drug resistance(14–18). However, all genetic systems developed so far for studying unstable genotypes rely on reporter genes, and thus investigate only one genetic locus, and only one type of genetic change.

Unbiased next-generation sequencing based approaches could give a more global view, allowing us to understand the rules that govern unstable genotypes at a genome-wide scale. However, genetic changes with high rates of reversion tend to remain subclonal(19–21), and it is challenging to distinguish most types of low-frequency mutations from sequencing errors(22), especially in complex genomes with large amount of repetitive DNA or *de-novo* duplicated genes. Thus, fast growing organisms with relatively small and simple genomes are particularly well suited for determining if transient mutations exist, for the genome-wide characterization of such mutations, and for identification of the underlying mechanisms.

## Results

### Microhomology mediated tandem duplications in specific genes caused reversible phenotypes in *S. pombe*

To discover novel transient drug resistance mechanisms in a eukaryote we performed a genetic screen in the fission yeast *S. pombe* for spontaneous mutants that are reversibly resistant to rapamycin plus caffeine (caffeine is required for rapamycin to inhibit growth in *S. pombe*(23)) **(Fig. 1A)**. We plated 10^7^ cells from each of two independent wild-type strains to YE5S+rapamycin+caffeine plates, and obtained 173 drug resistant colonies, 14 (7%) of which exhibited reversible drug resistance following serial passage in no-drug media **(Fig. 1B,C)**. In contrast, resistance for deletion mutants such as *gaf1Δ*(24) is irreversible suggesting, the existence of a novel type of genetic or epigenetic alteration allowing for reversible drug resistance in the newly isolated strains **(Fig. 1B,C)**.

**Figure 1.**
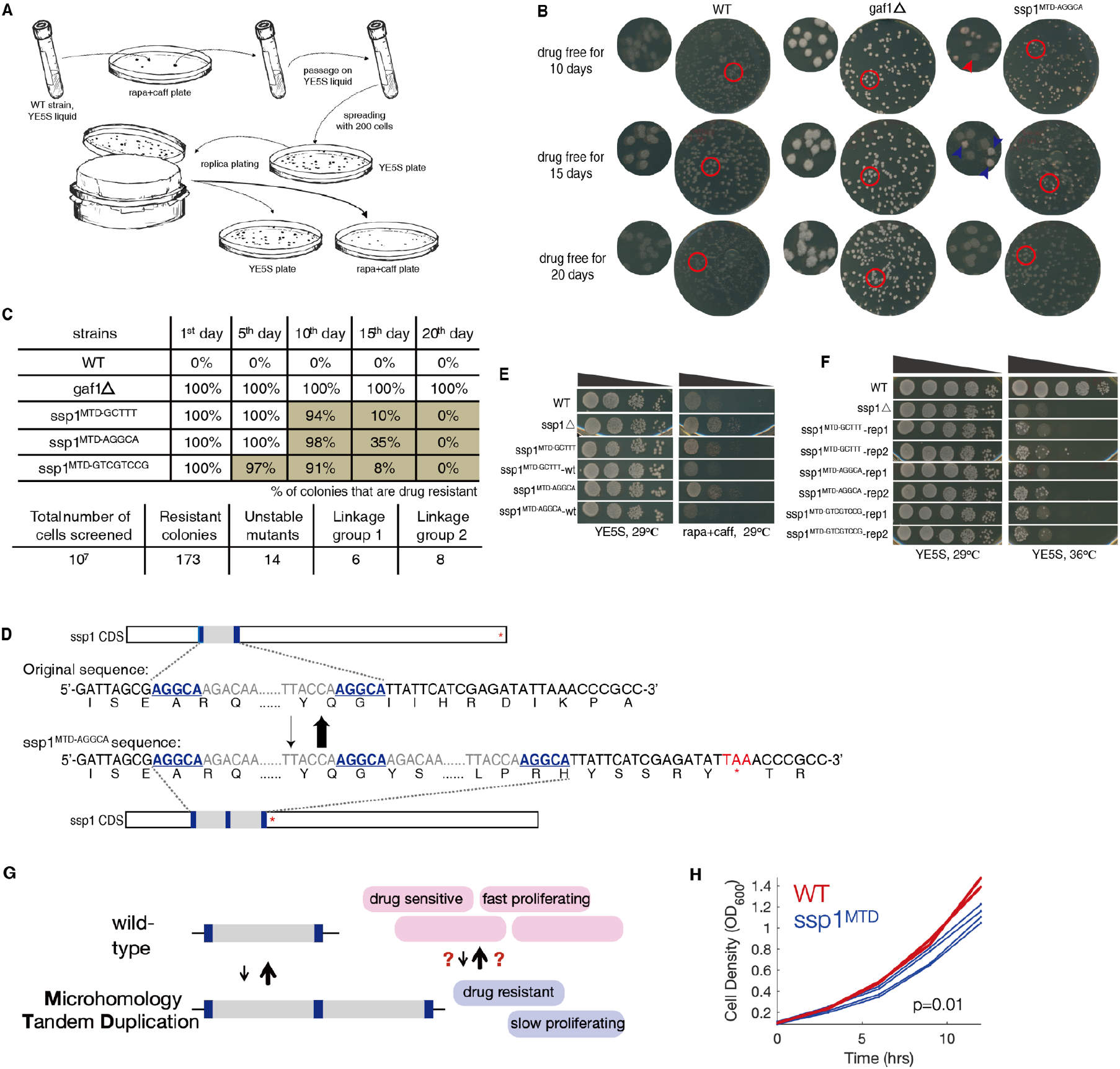
Screen for mutants with unstable inherited resistance by rapamycin plus caffeine and identify highly reversible mutations in *ssp1*. **(A)** Procedure to screen mutants with unstable rapa+caff resistance using sensitive wild-type strains in S. *p*ombe. **(B)** Unstable phenotype for one of screened mutants on rapa+caff plates after replica plating. *gaf1Δ* as positive control shows strong and stable resistance. The days represent for incubation time on drug free condition allowing the growth of resistance degenerated progeny. The red arrows point to sensitive progenies, while the blue to resistant ones. **(C)** Dynamics of reversion among identified reversibly-drug-resistant colonies. **(D)** Identification of tandem segment duplication in *ssp1* for drug resistance progenies by whole genome sequencing and reconfirmation by locus-specific PCR/Sanger sequencing. Underlined and bold bases stand for the microhomology pair. The pre-matured stop codon is marked with red. **(E)** *ssp1* inactivation caused rapamycin resistance and the replacement of ssp1^MTD^ sequence to wt-ssp1 rescue the drug resistance to wt level. **(F)** Heat-resistant isolates are frequently obtained in ssp1^MTD^ strains. **(G)** A cartoon of reversible MTDs that cause drug resistance and a proliferation defect. **(H)** Growth curves of wild-type (red, two replicates) and ssp1MTD^AGGCA^ (blue, four replicates).

We used genetic linkage mapping and whole-genome sequencing to identify the molecular basis of reversible rapamycin+caffeine resistance. We identified two linkage groups **(Fig. S1A)**; we could not identify any common mutations in the first linkage group, suggesting an epigenetic or non-nuclear genetic mutation, or an inheritable variation that remains to be detected. In contrast, all eight strains in the second linkage group contained novel tandem duplications in the gene *ssp1*, a Ca^2+^/calmodulin-dependent protein kinase (human ortholog: CAMKK1/2) which negatively regulates TORC1 signaling, the pathway inhibited by rapamycin, suggesting that mutations in *ssp1* were causal for drug resistance(25).

The *ssp1* linkage group contained three insertion alleles, all of which were tandem duplications of a short DNA segment (55/68/92 bps in length) and had 5-8 bp of identical sequence (MicroHomology Pairs, MHPs) at each end **(Fig. 1D, Fig. S1B, Table S7)**. We postulate these Microhomology-mediated Tandem Duplications (MTDs)(26–28) are important for de-novo generation of reversible mutations.

All three MTDs resulted in frameshifts and inactivation of *ssp1*. A similar level of drug resistance was found in the *ssp1Δ*, and replacement of the MTD alleles by transformation with wild-type *ssp1* restored sensitivity **(Fig. 1E)**. Sanger sequencing showed that all 16 randomly selected drug-sensitive revertants of the MTD alleles had the wild-type *ssp1* sequence. Finally, *ssp1Δ* and *ssp1*^MTD^ strains are temperature sensitive. Spontaneous drug-sensitive non-ts revertants were frequently recovered for all the *ssp1*^*MTD*^ alleles at a frequency of roughly 1/10,000 cells, but not for the *ssp1* deletion **(Fig. 1F)**. The frequency of revertants is thus 100x higher than the forward MTD frequency (8/10^7^), and MTDs in *ssp1* are causal for reversible temperature sensitivity and drug resistance.

Supporting the notion that MTDs may not be specific to rapamycin/caffeine treatment and/or the target gene *ssp1*, in an unrelated genetic screen for suppressors of the slow growth defect of *cnp1-H100M*, a point mutation in the centromere-specific histone gene, we identified MTDs in the transcription repressor genes *yox1* and *lsk1* **(Fig. S1B, S2, Table S7)**. These MTDs increase fitness in the *cnp1-H100M* background and therefore, unlike *ssp1*^*MTDs*^, revertants do not increase in abundance in the mutant background. However, in the *ssp1*^*wt*^ background, these MTDs are deleterious and revertants accumulate **(Fig. S1, S2)**. Thus, MTDs are not gene-specific and likely occur throughout the genome.

### 10,000x whole-genome sequencing identified thousands of subclonal MTDs within a clonal population

Based on the scale of the initial genetic screen, and assuming drug resistance is not induced by rapamycin, the frequency of cells with any protein-inactivating MTD in *ssp1* in an exponentially growing non-selected wild-type population is estimated approximately 8×10^−5^. This suggests that a clonal, presumed “isogenic” population contains a wide variety of subclonal MTDs at multiple loci throughout the genome. The frequency of any single MTD will depend on the rate of MTD formation, the rate of reversion and the fitness cost of the MTD (19–21). The fitness defect imposed by the MTD can be due to altered gene expression or protein function, or from the fitness cost of ∼0.025% per kb of additional DNA (29, 30).

To identify the *cis*-encoded determinants of MTD frequency we developed a computational pipeline for detecting subclonal MTDs in high-coverage Illumina sequencing data (see Methods for details). This method first identifies all MH Pairs (MHPs) in a DNA segment or genome and generates ‘signatures’ for sequences that would be created by each possible MTD. It then identifies sequencing reads that match these signatures, and thus provides experimental support for the existence of a particular MTD within the population **(Fig. 2A)**. This method is capable of identifying subclonal MTDs present at very low frequencies in the population.

**Figure 2.**
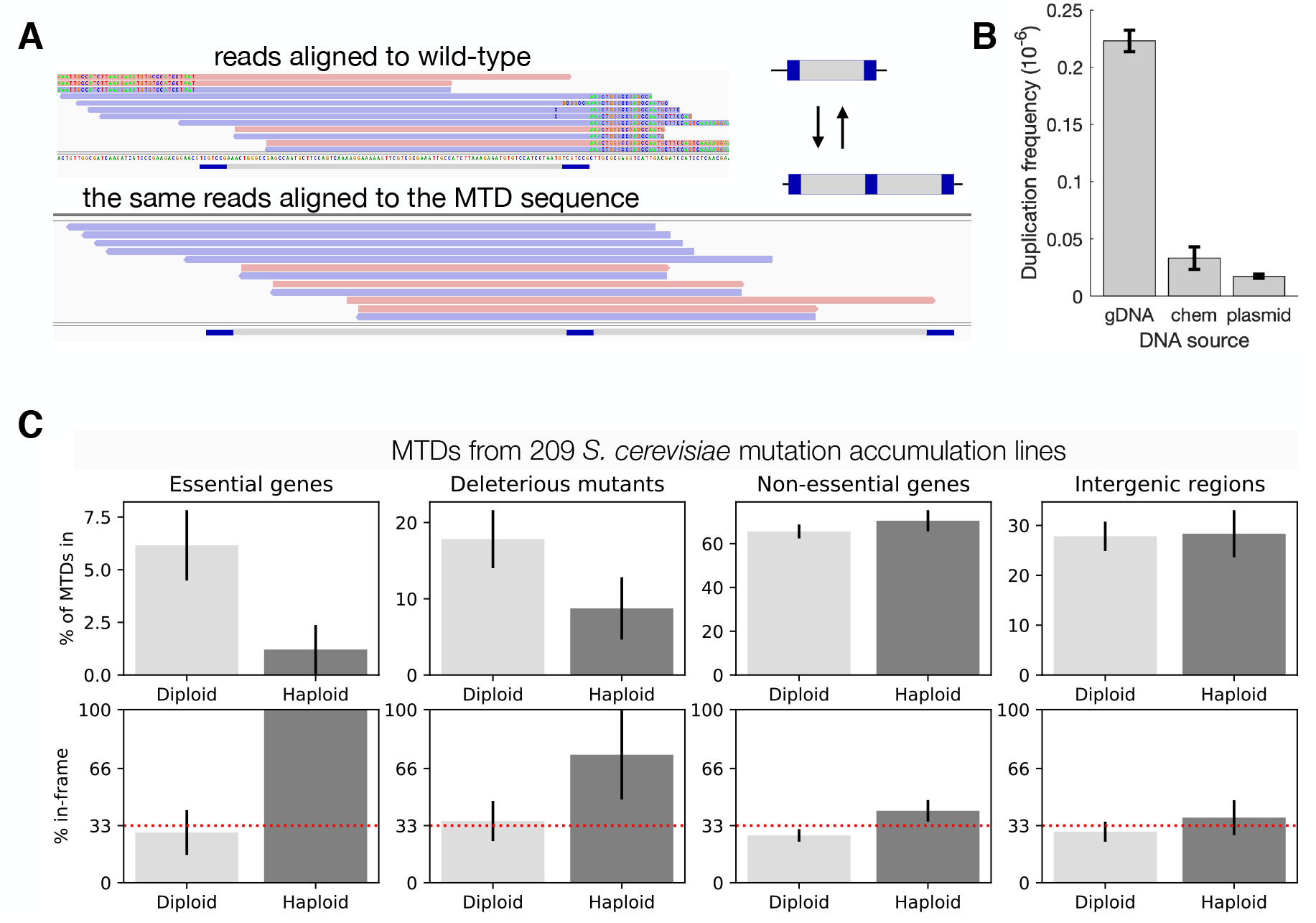
Identification and verification of subclonal MTDs from ultra-deep sequencing data. **(A)** The computational pipeline finds all sequencing reads that whose ends do not match the reference genome, and checks if the reads instead match the sequence that would exist due to an MTD. Shown are reads identified in the pipeline, aligned to either the reference genome (top) or to a synthetic genome with the MTD (bottom). Red and blue mark reads that map to opposite strands. The MHPairs are shown in dark blue, and positions in each read that do not match the reference are colored according to the base in the read. **(B)** The average frequency of sequencing reads that support each MTD in ssp1 from 10^6^ coverage sequencing of the gene from *S. pombe*, from a plasmid-borne *ssp1* in E. coli, or from a chemically synthesized fragment of the *ssp1* gene. Error bars are standard error of the mean across replicates. **(C)** The genomic locations of 314 MTDs that occur in only one single mutation accumulation line, reanalyzed raw sequencing data from (32); error bars are standard deviation from bootstrapping. The top row shows the percent of MTDs found in essential genes, in genes whose haploid deletion mutant is viable but shows a fitness defect (33), in all non-essential genes, or in intergenic regions, separately for haploid and diploid M.A. lines. The bottom shows the percent of MTDs in each category that create a tadem duplication that has a length divisible by three, and therefore would not disrupt the reading frame. The red dashed line shows the random expectation (1/3^rd^ of MTDs).

To determine if subclonal MTDs captured by sequencing represent the true genetic variation, or are technical artifacts (31) we performed two orthogonal tests. In the first, we tested if MTDs are specific to genomic DNA, or also exist in chemically synthesized DNA. We performed 10^5^x - 10^6^x coverage sequencing of *ssp1* DNA fragments PCR-amplified from genomic DNA, from a cloned copy of the gene in a plasmid in *E. coli*, or chemically synthesized 150nt and 500nt fragments of the gene, as well as direct sequencing of chemically synthesized short DNA fragment and plasmid-borne fragment without PCR amplification. We observed far more MTDs in the *pombe* genomic DNA than in the chemically synthesized or plasmid borne controls **(Fig. 2B, Fig. S3)**, suggesting that MTDs are largely not caused by PCR or an artifact of Illumina sequencing. It is unclear why the plasmid-borne copy of *ssp1* has fewer MTDs than the chemically synthesized DNA but it raises the possibility that MTDs may be more frequent in eukaryotes **(see also Fig. 2D, S6, S7, S8)**. We detect MTDs in the *E. coli* genome 1/20^th^ as often as in *S. pombe* and 1/60^th^ as often as in *S. cerevisiae* **(Fig. S9)**.

As a second test, we hypothesized that most MTDs in essential genes should be deleterious and recessive. We therefore analyzed raw sequencing data from 209 *S. cerevisiae* haploid and diploid mutation accumulation(MA) lines(32) and identified all MTDs that occur only one MA line. In haploids, MTDs were depleted in both essential genes and non-essential genes whose deletion causes a fitness defect **(Fig. 2C)**. In addition, the MTDs that did occur were more likely to maintain the correct reading frame; the single MTD in an essential gene in a haploid was subclonal, maintained the correct reading frame, and was just 112bp from the 3’ end of CCT7, a 1652bp gene. In contrast, there was no such reading-frame bias in diploids, non-essential genes, or intergenic regions **(Fig. 2C)**. Therefore, rare subclonal MTDs identified by ultra-deep sequencing are likely real biological events mostly not experimental artifacts.

To assess the prevalence of MTDs and to identify the sequence-based rules that determine the probability of formation of each tandem duplication, we grew a single diploid fission yeast cell up to ∼10^8^ cells (25 generations) and performed whole-genome sequencing to an average coverage of 10,000x. The diploidy relaxed selection, allowing mutations to accumulate throughout the *S. pombe* genome.

With 10,000x genome sequencing, we identified 5968 (0.02%) MHPs in which one or more sequencing reads supported an MTD. We observed zero MTDs in most genes, likely due to under-sampling **(Fig. S4)**. However, 20 genes contained more than ten different MTDs in a single ‘clonal’ population **(Fig. 2A)**. To understand this heterogeneity across the genome we used a logistic regression machine-learning model to predict the probability of duplication at each MHP. MH length, GC content, inter-MH distance, measured nucleosome occupancy, transcription level, and a local clustering on the scale of 100nt, were able to predict which MHPs give rise to duplications with an AUC of 0.9 with 10-fold cross validation **(Fig. 2B,C, S5, Table S5)**. We note that the peak at 150nt inter MH spacing is independent of read length, was not found in *E. coli* or in mitochondrial DNA, and varies between haploid and diploid **(Fig. S5, S6, S7, S8)**. This analysis revealed properties of MHPs significantly affect the likelihood of MTD formation; for example, and consistent with previous work in *E. coli* (34), long GC-rich MH Pair is 1000x more likely to generate a tandem duplication than a short AT-rich one.

While MHPs are spread roughly uniformly throughout the genome **(Fig. 2D, red)**, we observed both hot-spots, in which MH-mediated generation of tandem duplications are common, and cold-spots, in which they are rare **(Fig. 2E)**. Local differences in MHPair density can only explain some of the hotspots, while our logistic regression model explains the vast majority, suggesting that hotspots with frequent formation of tandem duplications are mostly determined by the local DNA sequence features, in addition to microhomologies. The consequence is that duplications are more than 10x more likely to occur in some genes than others, and this variation is correctly predicted by our model **(Fig 2F)**. We detected no MTDs in *ura4*, which has a score of 52, placing it in the bottom third of genes **(Table S4, Fig. S10)**, and providing a possible explanation why MTDs have not been noticed in 5-FOA based screens of mutations in *ura4*(35). Our results also emphasize that high-coverage sequencing is necessary to identify sufficient numbers of MTDs; one billion reads would be required to identify half of the 25 million possible MTDs in the *S. pombe* genome **(Fig. S4)**.

We identified three different subclonal MTDs in the SAGA complex histone acetyltransferase catalytic subunit *gcn5*, placing *gcn5* in the top 5% of genes for both observed and predicted MTDs, suggesting that MTDs in *gcn5* should be found frequently in a genetic screen. Indeed, examination of 16 previously identified(36) suppressors of *htb1*^*G52D*^ identified MTDs in *gcn5*, as well as in *ubp8*, where we also observed an MTD in our high-coverage sequencing data **(Fig. S1B)**. These results suggest that MTDs arise in most genes at a high enough frequency within populations in order to be the raw material on which natural selection acts.

### Replication slippage modulates the rate of MTD reversion at *ssp1*

Having established that local *cis*-encoded features determine the frequency with which tandem duplications arise from microhomology-pairs, we next sought to identify the *trans*-genes that affect MTD process. *ssp1*^*MTD*^ alleles fail to grow at 36°C, and their reversion back to wild-type suppresses the temperature sensitivity, providing way to measure the effects of mutations on reversion frequency. We screened a panel of 364 strains with mutations in DNA replication, repair, recombination or chromatin organization genes for mutants that affect the rate of *ssp1*^*MTD*^ reversion back to wild-type **(Table S6)**, and found three mutants that significantly increased and eight that significantly decreased the frequency of *ssp1*^*WT*^ revertants **(Fig. 4A,B,C)**.

**Figure 3.**
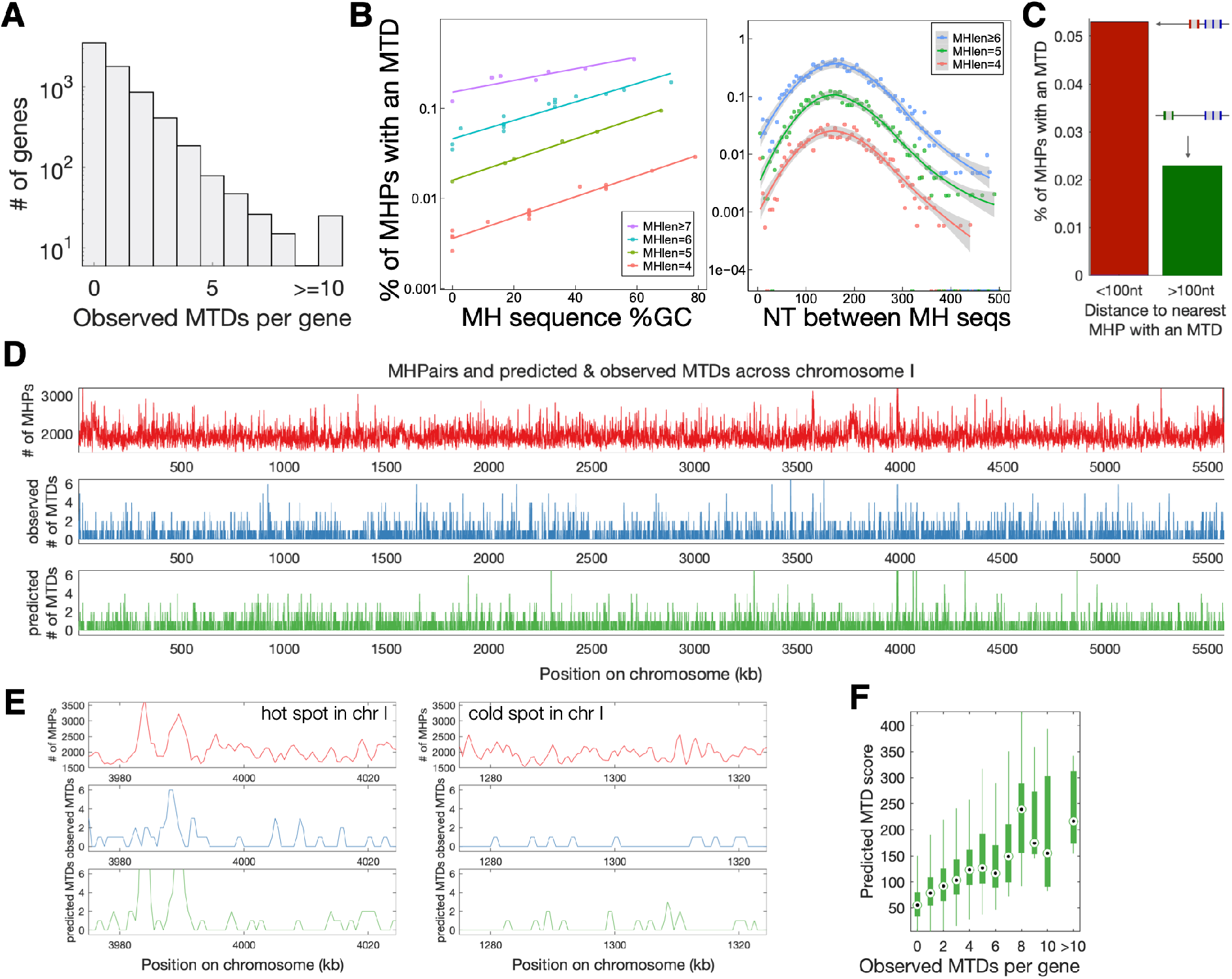
Identification of the *cis-*determinants of MTD through ultra-deep sequencing and identification of subclonal duplications. **(A)** A histogram of the number of MTDs found in each gene from 10,000x whole-genome sequencing. **(B)** The 25 million MHPs in the genome were binned in groups of 10,000 with the same MH sequence length and similar GC content (left) or inter-MHPair distance (right), and the % of MHPs in each group with an observed MTD was calculated. A logistic regression model was trained with 10-fold cross-validation to predict the probability of observing an MTD at each MHPair. **(C)** The distance from each MHP to the nearest MHP with an MTD was calculated, and the % of MHPs with an MTD was calculated for MHPs less than (red) or farther than (green) 100nt from the closest MHP. **(D)** For each 1kb window in the genome, shown are the number of MHPairs (red), the number of observed MTDs (blue), the predicted number of MTDs from the logistic regression model (green). **(E)** An example cold spot (0.2MTDs/kb) and hot spot (0.7 MTDs/kb) in chromosome I. The cold spot has fewer MTDs after taking into account the number of MHPs, (Fisher’s exact test, p=2.76e-09, odds ratio = 3.843). **(F)** The sum of scores from the logistic regression model for each MHP in each gene, with the genes grouped by the observed number of MTDs in the 10k coverage data.

**Figure 4.**
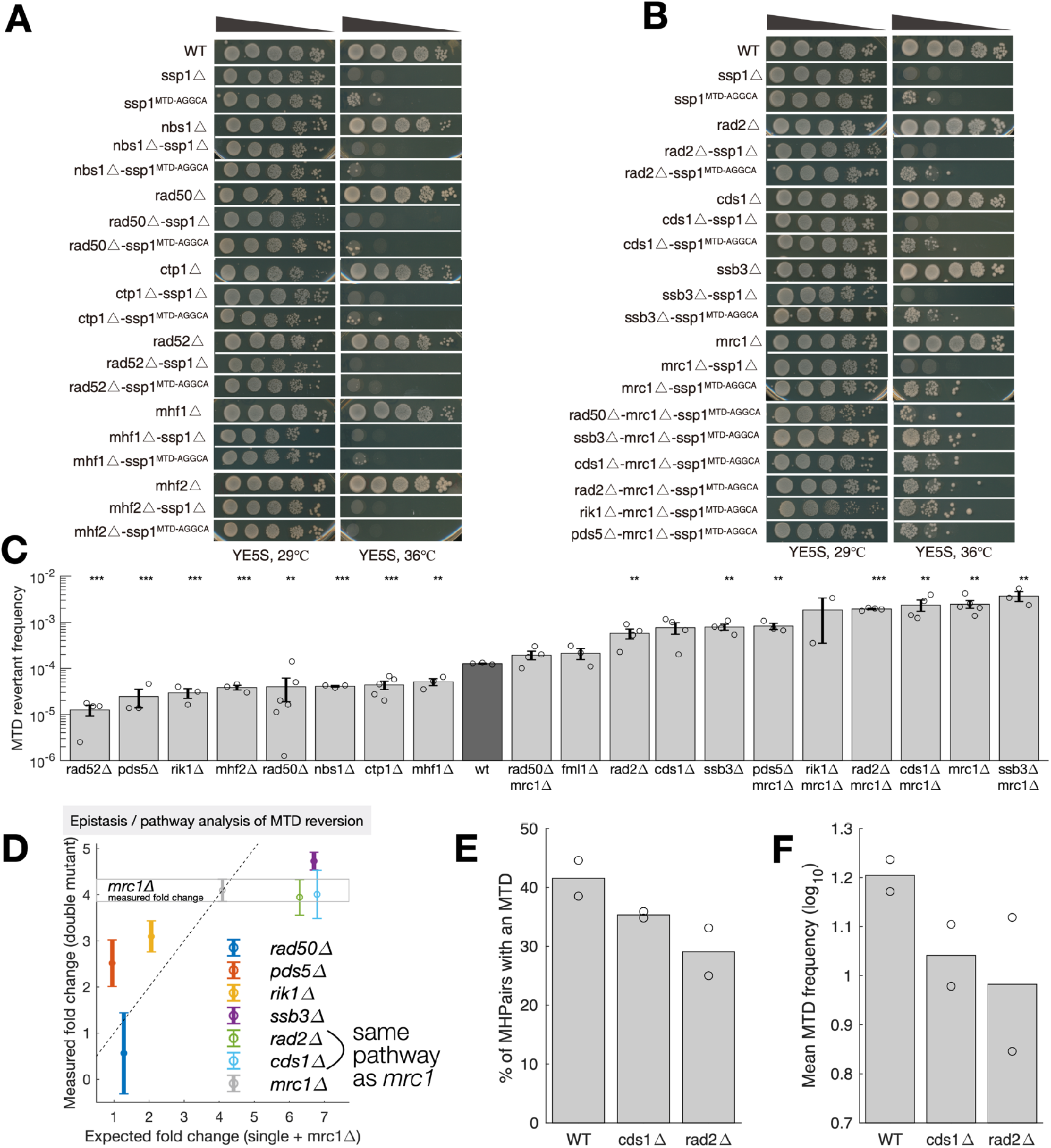
A genetic screen to identify the regulators of MTD reversion. **(A,B)**. Surveyed mutants showed reduced ssp1^MTD^ reversion frequency represented by TS recovery phenotype. The non-TS phenotype of single mutation and *ssp1Δ* alone or combined with other mutants retained severe temperature sensitive phenotype at 36°C should be established. The number of TS revertants under 36°C indicate the reversion frequency of ssp1^MTD^. The initial gradient for spotting assay was 10^5^ cells, and diluted with tenfold gradient (cell number: 10^5^, 10^4^, 10^3^, 10^2^, 10^1^). **(C)**. Quantification of ssp1^MTD^ reversion frequency in mutants (n>=3 biological repeats, error bars are s.e.m., *** = p<0.001, **=p<0.01, *=p<0.05 t-test compared to wt). **(D)** Two colonies of WT and two of each mutant were picked and SPCC1235.01 amplified by PCR and sequenced to 10^6^ coverage. Show is the average across the two replicates of the MTD frequency at each of the 3002 MHPs. **(E)** The % of MHPairs with one or more reads in support of an MTD in SPCC1235.01. **(F)** For all MHPairs with an MTD, the frequency of reads supporting that MTD per 10^6^ reads that map to that MHPair.

Replication fork collapse is a major source of double stranded breaks (DSBs), and the ensuing Homologous Recombination (HR)-related restarting process is error-prone and is known to generate microhomology flanked insertions and deletions via replication slippage, a process in which when replication resumes for a stalled or collapsed fork, the unwound nascent strand may anneal with a homologous segment on the template, either at the vicinity (37, 38) or at a distance(34) of the paused site, with ensuing replication on non-continuous template. Inactivation of Rad50, Rad52 or Ctp1 results in decreased replication slippage, and decreased MTD reversion (37, 39)**(Fig. 4A,B,C)**. Deletions of *mhf1* and *mhf2*, two subunits of the FANCM-MHF complex, which is involved in the stabilization and remodeling of blocked replication forks, also decreased the frequency of MTD revertants. It is therefore likely that replication slippage during HR-mediated fork recovery causes reversion of MTDs in these mutants and could be one contributing factor in wild type background.

Replication stresses activate a checkpoint that promotes DNA repair and recovery of stalled or collapsed replication forks, and delays entry into mitosis(40, 41). The inactivation of replication checkpoint kinase *cds1* or its regulator *mrc1* may thus result in a failure to restore the replication fork, causing increased genome instability and MTD reversion. The replication checkpoint would thus be required for the stability of MTDs. Consistently, we found that deletion of the DNA damage checkpoint kinase *cds1* or its regulator *mrc1* increased the frequency of *ssp1*^*W*T^ revertants. Deletion of the single-stranded DNA binding A (RPA) subunit *ssb3* (RPA3/RFA3) or the multifunctional 5’-flap endonuclease *rad2* also increased the frequency of revertants **(Fig. 4C)**.

Many genes identified in the screen are multifunctional, and play roles in both replication and repair. We therefore performed quantitative epistasis analysis to determine the relation between six of the identified genes and the Mediator of the Replication Checkpoint, *mrc1*, which interacts with and stabilizes Pol2 at stalled replication forks. In addition to the checkpoint activator *cds1*, deletion of *rad2* had no effect in an *mrc1Δ* background, suggesting that all three of these genes act in the same pathway **(Fig 4D)**. In contrast, deletion of *ssb3* increased the frequency of revertants in both wild-type and *mrc1Δ* backgrounds, and deletion of *pds5* or *rik1* decreased the frequency of revertants in both wild-type and *mrc1Δ* backgrounds, though not to the extent expected for genetic independence, suggesting partial epistasis. In contrast, the effects of *rad50* deletion were completely independent of *mrc1* **(Fig 4D)**.

While the observed numbers of MTDs in ultra-deep sequencing experiments are a function of both duplication and reversion rates, and all of the above genes may play a role in both processes, the above results suggested that, due to increased reversion rates, the number and frequency of MTDs would be reduced in *cds1Δ* and *rad2Δ* strains. To test this we performed 10^6^x coverage sequencing of the hotspot gene *SPCC1235*.*01*. We observe MTDs at fewer MHPairs, and an overall decrease in the number of MTDs in both mutants **(Fig. 4E,F)**.

### Half of insertions and tandem duplications in natural isolates are MH-mediated

It was baffling that MTDs are prevalent within populations, and that the first theoretical proposal for microhomology-mediated processes in the generation of tandem duplications is twenty years old(5), yet, relatively little is known about the forward process, and even less about the reversion, suggesting that these events are not often encountered or identified as such. To better understand the dynamics of MTDs within a population we used a simple model of neutral mutations within a growing population that takes into account both forward and reverse mutation rates and began with 100% of individuals as wild-type (see Methods). The mutant frequency always increases, and over short timescales **(Fig. 5A**, left) increasing the reverse rate from being equal to the forward mutation rate (grey) to being 10,000 times higher (yellow) has little effect.

**Figure 5.**
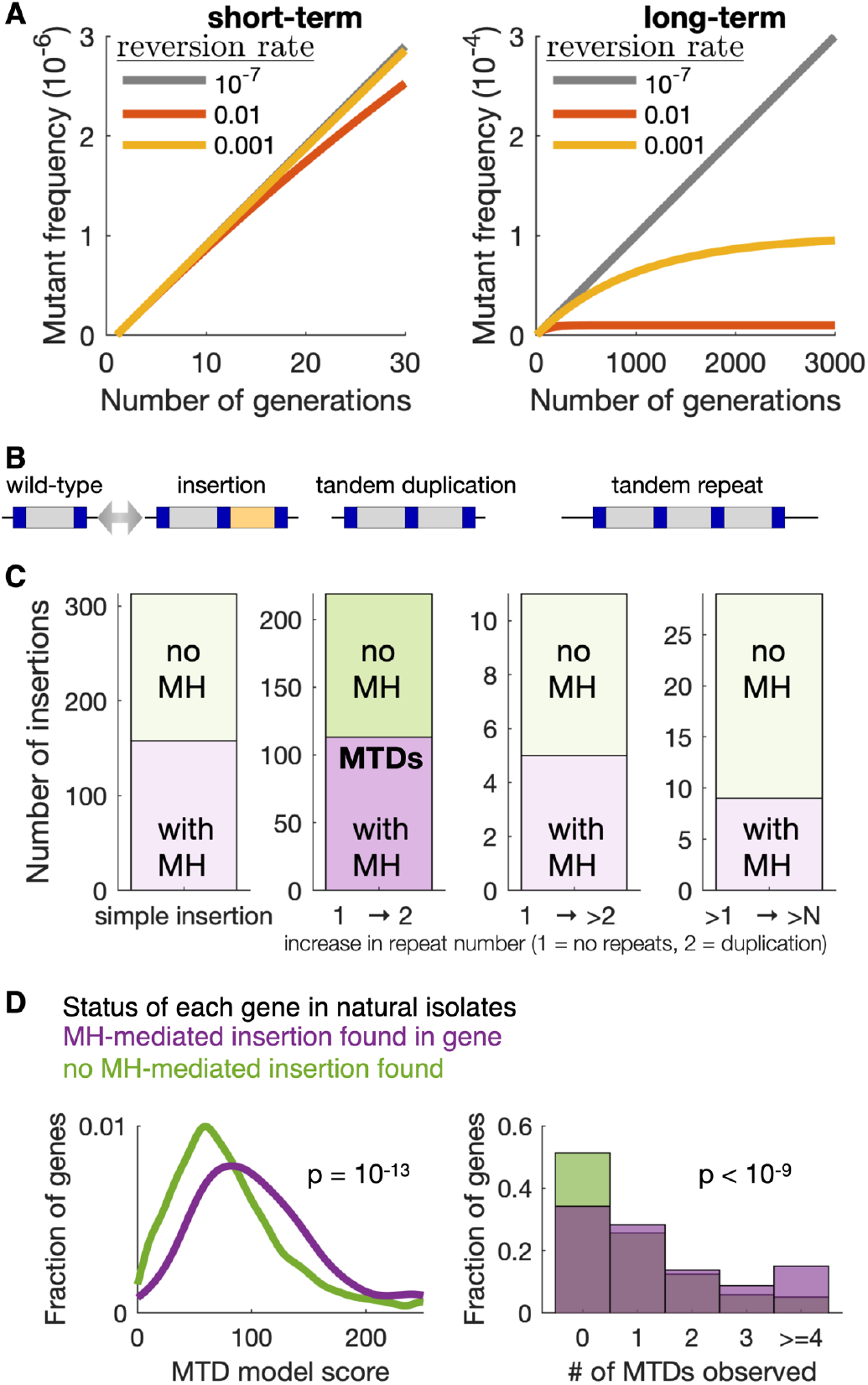
MTDs remain subclonal due to high reversion rates, yet half of insertions and de-novo tandem duplications in natural populations arise at microhomology sequence pairs. **(A)** Simulations showing the frequency of a neutral mutation (forward mutation rate = 10^−7^) within a growing population at three different reversion rates (colors). Left and right show the same simulates at different timescales, with the effect of reversion only apparent at long timescales. **(B)** A cartoon showing three possible types of microhomlogy mediated insertions: simple insertion, tandem duplication, and higher copy repeat. **(C)** Quantification of all insertions of at least 10bp fixed in any of the 57 natural *S. pombe* isolates that represent most of the genetic diversity within the species, relative to the reference genome. Insertions were classified according the presence (purple) or absence (green) of exact microhomology pairs on either side of the insert, and to the type of insert. There are 113 MTDs in wild *pombe* strains (second column). The right-most column (>1x -> >Nx) refers to the expansion of repeats present in the reference genome. **(D)** Distributions of the predicted MTD score from the logistic regression model (left) and the number of experimentally observed subclonal MTDs (right) for genes with one or more microhomology-mediated insertions (purple) or for genes with no MH-mediated insertions (green) in any of the natural isolates. p-values are from a Mann–Whitney U test.

Over longer timescales, high reversion rates cause the mutant frequency to plateau and remain subclonal **(Fig. 5A**, right), reducing the fraction of neutral MTDs within a population. However, in spite of the high reversion rate, both drift and selection enable fixation of MTDs within a population. To identify fixed microhomology mediated insertions we searched the genome sequences of 57 wild *S. pombe* isolates (42), and found that 50% of insertions larger than 10bp involve microhomology repeats **(Fig 5B,C)**. Among these were 158 microhomology mediated insertions that did not contain an obvious duplication, and 113 MTDs with a microhomology mediated tandem duplication.

To test if the propensity of MTD formation within the lab strain is predictive of extant sequence variation observed in natural isolates, we tested if the MTD score for each gene predicts the likelihood of microhomology mediated insertions in that gene. We found that genes with microhomology mediated insertions in natural isolates tend to have higher predicted MTD scores, and more experimentally observed MTDs **(Fig. 5D)**, suggesting that the local features that affect MTD formation in the lab also shape evolution in nature.

### MHPs with longer MH sequences are more likely to generate MTDs that maintain the correct reading frame

We found that MHPs with longer MH sequences are more likely to form MTDs. If the high propensity to generate MTDs has shaped the *S. pombe* genome, any signature of selection should be stronger at MHPs with longer MH sequences, and should also be stronger in essential genes vs non-essential genes. We therefore divided the 25 million MHPs with an MH length of 4-25nt and an inter-MH distance of 3-500nt into those fully contained within in intergenic regions, or fully contained within essential or non-essential genes, and, split them by MH sequence length. Specifically in coding sequences, MHPs at which an MTD would not disrupt the reading frame are more common than expected by chance, and this enrichment is higher in essential genes, and at longer MH sequences **(Fig 6)**. At the same time, MHPs within genes are more common that expected by chance **(Fig. S12)**. Therefore, natural selection has acted to decrease the number of MHPs that would create potentially deleterious MTDs, and this selection is weaker for MHPs that would create an in-frame MTD.

**Figure 6.**
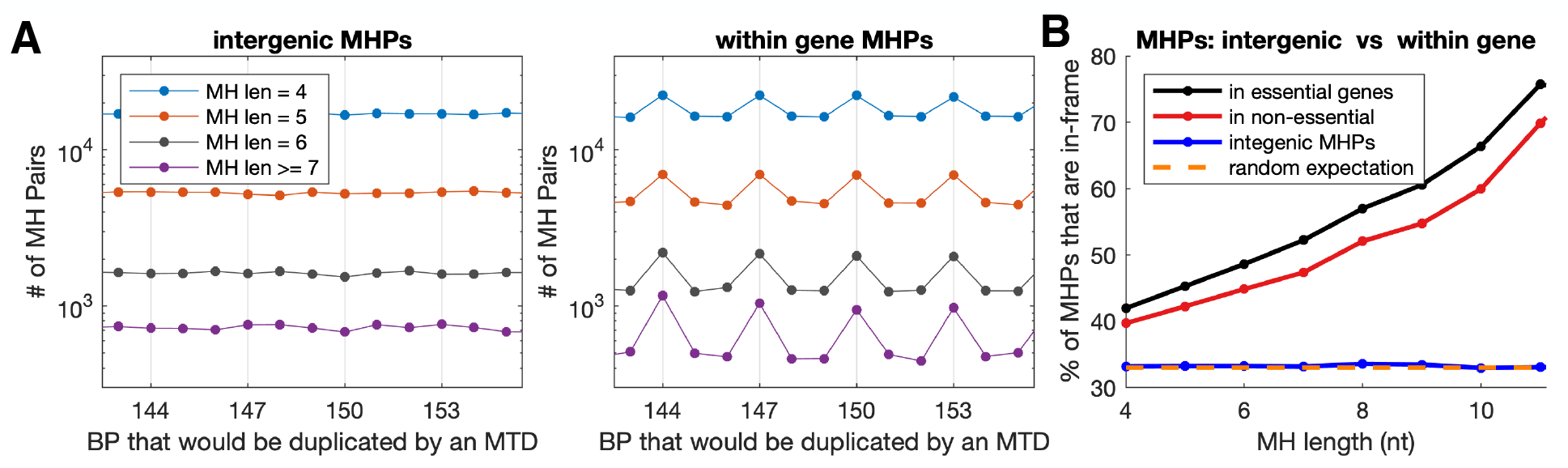
Frame-shifting MHPs are depleted from essential genes in an MH-sequence-length dependent manner. **(A)** The number of MHPs in the *S. pombe* genome with different MH sequence lengths (colors) for which an MTD would generate varying insert sizes (x-axis). X-axis grid lines mark MTDs with insertion sizes divisible by three. Left shows MHPs that are intergenic, and right MHPs that are fully contained with a coding sequence of a gene. **(B)** The % of MHPs with lengths evenly divisible by three (y-axis) for each MH sequence length (x-axis) that are found in intergenic regions (blue), fully contained within essential genes (black) or within non-essential genes (red). Random expectation is that 1/3^rd^ of MHPs will have an insert size evenly divisible by three (orange).

Taken together, our results demonstrate that MTDs occur frequently and broadly throughout the genome within a clonal population. This indicates that high levels of subclonal genetic divergence are prevalent but are under-detected using conventional sequencing approaches that tend to disfavor the detection of low abundance subclonal variants. As many MTDs create large insertions, they are more likely to be deleterious. Nonetheless, MTDs provide plasticity to the genome and its functionality, for example, by allowing cells to become drug resistant, while allowing the resistant cell lineage to revert back to wild-type and regain high fitness once the drug is removed. Selection can act on this genetic diversity for its reversibility or by using the tandem duplications as the initial step for the generation of higher copy number repeats, which are evolutionarily fixed in extant genomes and traditionally regarded as a major source of genome divergence. While previous work has shown that pre-existing repeats undergo rapidly reversible changes, the sequence-encoded rules regulating the birth and death of such sequences were less studied(34). This work reveals that numerous sites throughout the genome have the potential of evolving into such repetitive elements. Furthermore, MH-sequence length dependent depletion of frame-shifting MHPs in essential genes shows that natural selection has shaped the genome to avoid MHPs that would frequently generate deleterious MTDs. Finally, much in the same way as repetitive DNA may have been positively selected for as a regulatory element to maintain reversible genetic diversity(7, 8), the large number of MHPs that would result in in-frame MTDs raises the possibility that some genes may maintain MHPs to generation functional genetic diversity, creating a dynamic protein-coding genome.

## Discussion

Why haven’t MTDs been identified more frequently in genetic screens and mutation accumulation assays? There are several possible. Mutation callers are ineffective in detecting long insertions from short reads: on simulated data, both Mutect2 and HaplotypeCaller often fail to detect tandem duplications longer than 85bp (data not shown). Also, MTDs are often identified as insertions, but not specifically as MTDs **(Fig. S1B, S11)**, suggesting the need for computational tools for identifying MTDs. URA4 and URA3 have relatively few MHPs **(Fig. S10)**. In many 5-FOA-based mutation spectra papers, only substitutions were analyzed in detail, as indels do not occur with high enough frequency to generate good statistics(43). Due to the high rate of reversion, it is likely that tandem duplications in URA4 would yield URA+ colonies when restruck onto -URA plates; this high reversion rate may lead to MTD-containing colonies being discarded in many different types of genetic screens. In our re-analysis of *S. cerevisiae* mutation accumulation lines, almost all de-novo MTDs were subclonal **(Fig. S11B)**. With new computational tools for identifying MTDs, plus third-generation sequencing platforms with improved ability to detect long indels, it is likely that MTDs will be implicated in more phenotypes.

Unbiased genome-scale approaches have been very informative for the mechanisms that generate point mutations in both wild-type and mutant cells(44). It is clear that multiple molecular mechanisms can give rise to tandem duplications, in microhomogy dependent and independent manners, and the mechanisms may differ in mutants and between species. In plant mitochondria genomes, longer MH sequences are associated with longer tandem duplications, suggesting that microhomology is involved in the generation of tandem duplications, likely via microhomology-mediated repairing of DSBs (45, 46) or slippage strand replication (47). In contrast, tandem duplications in the rice nuclear genome tend to have no or shorter MH sequences, suggesting that in in the rice nuclear genome, tandem duplications likely form via patch-mediated DSB creation followed by NHEJ (48). In *E. coli*, the lagging-strand processing activity of Pol I is required for stress-induced MH-mediated amplification of 7–32kb segments(34). However, simple models cannot account for all features observed across studies, and it is clear that multiple mechanisms play a role (49–51).

All current methods of measuring microhomology mediated duplications and deletions impose artificial length scales whereas genetic screens require the entire gene or a specific region be duplicated. Thus, while microhomologies are associated with deletions on the 500bp-1kb range in E. coli(52), and with unstable amplifications of 7-37kb(16) and that selection for increased gene expression often enriches for ∼12bp MH-mediated amplification of ∼10kb (53), these length scales are determined by the locations of MH sequences in the particular genomic region required to be duplicated in the genetic screen, and the size of the region required to be duplicated or amplified.

Genome-wide sequencing based approaches are less biased, but still not bias-free. We limit the length-scale to 500bp and set lower and upper bounds of 3bp and 25bp for the MH sequences. While the MTD frequency, relative to the number of 1bp or 2bp MH sequence pairs in the genome is likely to be low, the number of 2bp and 1bp MH sequences is high. Preferential flanking of tandem duplications by 2bp and even 1bp sequence identities have been reported(27, 28, 45, 54), suggesting that sub-clonal MTDs generated by short MH sequences may be common. While it is computationally intractable to apply the directed ‘signatures’ method presented here to search MH pairs separated by more than 500bp, this could be extended to >500bp if the search is limited to longer MH sequences, which are rare, but drive high rates of MTD formation(28, 34). Similarly, the method could be extended to broken microhomologies(55), but with an even higher computational cost. Computational approches using third-generation (Nanopore and PacBio) sequencing have the potential to provide a truly unbiased measure of duplication and deletion frequencies, as well as answer questions about how often amplifications are extrachromosomal vs intrachromosomal (17, 56), and tandem vs inverted amplifications (57). Verification of these algorithms will take some work, as nanopore and Pac-Bio sometimes give different results when sequencing tandem repeats(58).

Because different molecular mechanisms have different sequence requirements, replication strand biases, and length scales(34, 50), genome-wide unbiased methods are necessary to understand the relative contribution of each mechanism to MTD formation and collapse. As they can, in theory, measure events across any distance, from single bp to inter-chromosomal, and at an unlimited number of different loci, with variation in chromatin contexts, transcription, and other genome-architecture features, ultra-deep sequencing is likely the best way to quantitatively understand the various biological mechanisms that contribute to the dynamic genome.

## Acknowledgments

We thank Lilin Du, Aaron New, and Wenfeng Qian for insightful discussions and for comments on the manuscript. We thank Qi Zhou for assistance with rapamycin and caffeine resistance screen.

## Funding

L.B.C. was supported by the Peking-Tsinghua Center for Life Sciences. X.H. was supported by National 973 Plan for Basic Research Grant 2015CB910602 and National Natural Science Foundation of China Grant 31628012.

## Author contributions (CRedIT)

Author contributions (CRediT):
| AUTHORS | ROLE |
| --- | --- |
| X.H. | Conceptualization |
| L.D., S.C., Y.L., X.H., L.B.C. | Methodology |
| S.C., Y.L., & L.B.C. | Software |
| L.D., S.C., Y.L., & L.B.C. | Validation |
| L.D., S.C., Y.L., & L.B.C. | Formal analysis |
| L.D., S.C., Y.L., J.L. & L.B.C. | Investigation |
| X.H., F.L., Y.W. | Resources |
| L.D., S.C. Y.L. & L.B.C. | Data Curation |
| L.B.C. L.D. & X.H. | Writing |
| L.D., S.C. X.H. & L.B.C. | Visualization |
| L.B.C. & X.H. | Supervision |
| L.B.C. & X.H. | Project administration |
| L.B.C. & X.H. | Funding acquisition |

## Competing interests

Authors declare no competing interests.

## Data and materials availability

All processed data and code are available at https://github.com/carey-lab/MicroHomologyMediatedTandemDuplications and raw sequencing data at NCBI SRA BioProject accession PRJNA631756.

## Materials and Methods

### Strains

*S.pombe* strains used in this study are listed in Table S1. The deletion strains and GFP-tagging strain were originated from the genome-wide deletion library(59) or constructed by overlap PCR strategy and gene-specific homologous recombination using standard procedures(60).

### Cell Growth

Fission yeast cells were grown on YE5S liquid or solid medium (5S: supplemented with histidine, uracil, lysine, leucine, adenine), mated or sporulated on specific malt extract (ME) agar medium following standard procedures(61). For the preparation of rapamycin plus caffeine drug plate, 1000X stock solution of rapamycin(100μg/ml) was prepared by adding 100mg rapamycin to 1ml DMSO (100mg/ml) and diluting by 1000 folds. 1.942g power caffeine was dissolved in 60-80°C 20 ml sterile ddH_2_O and added into 1L YE5S medium to final concentration 10mM.

### Unstable drug-resistant mutants screen

A fresh single colony of wild-type cells was picked and grown to mid-log phase culture. Cultivated cells were then spread on YE5S agar plates containing 100ng/ml rapamycin and 10mM caffeine (hereafter called YE5S+drug plates) at the density of 1×10^5^ cells per plate and incubated at 29°C for 10 days. To test the stability of the drug resistance, each strain is grown continuously in YE5S liquid media in the absence of the drugs at 29°C by refreshing the culture with YE5S liquid media daily for up to 20 days. Every five days, cell samples were taken and spread to the YE5S plate at the density of 200 cells per plate. After a 3 day incubation at 29°C, each plate was replica plated to fresh YE5S and YE5S+drugs plates, respectively, incubated for two days at 29°C. Plates were visually examined for colonies that grow on YE5S but fail to grow on YE5S+drugs plates. The stability test was repeated at least two times for identified unstable drug-resistant strains. The *gaf1-d* mutant was used as the control for stable and robust drug resistance.

### Genetic linkage test

Identified unstable drug-resistant strains were backcrossed with wild-type cells or crossed with each other on the ME plate. After 24-48h sporulation at 29°C, tetrad-dissection was performed on the YE5S plate following the standard procedure(62). After 3 days incubation at 29°C, YE5S plates are replica plated to the YE5S+drug plate and incubated at 29°C for 2 days to identify drug-resistant colonies among the four progeny that originated from one ascus. The segregation pattern of the drug-resistant and drug-sensitive phenotypes is analyzed and used to determine the genetic linkage of the tested mutation alleles.

### Whole-genome sequencing and datasets analysis

Genomic DNA was extracted using phenol-chloroform, mechanically sheared to ∼200bp using ultrasonicator. Sheared genome DNA was used to build the library using NEBNext® Ultra™ DNA Library Prep Kit for Illumina® (E7370/7335, NEB) and Illumina sequenced by Ribobio in Wuhan, China.

To identify mutations in the genetic screens, adapter-trimmed FASTQ clean data were mapped to the ENSEMBL Fungi’s *S. pombe* genome version ASM294v2 with the BWA mem aligner (63)(version 0.7.17, with -M flag on). After removing PCR duplicates (64), BAM files were fed to the GATK’s HaplotypeCaller, base scores recalibrated using GATK’s BaseRecalibrator, and the recalibrated BAM files were input to the HaplotypeCaller to generate raw mutation callings(65), which were filtered and annotated using the ENSEMBL’s variant effect predictor (VEP, version 93.3)(66).

### Double mutant construction and MTD reversion regulators survey

Double mutants which combine MTD mutation at *ssp1* (*ssp1*^*MTD-AGGCA*^) and each deletion within the mutant panel were created by genetic crossing following the standard procedures (Roguev et al., 2018; Schuldiner et al., 2006) using a high throughput robotic apparatus (Peking university, F. Li lab. Protocol for high throughput manipulation is available upon request).

To assess MTD reversion rates semi-quantitatively, a single colony of each double mutant was used to inoculate 3ml YE5S liquid culture, incubated at 29°C overnight, refreshed by 1:10 dilution in 20ml YE5S liquid medium, and grown to mid-log phase. Serial 1:10 dilutions of the culture were prepared using fresh YE5S liquid medium, spotted on YE5S plates (57l per spot, corresponding to 10^5^ to 10 cell per spot), incubated for 4-5 days at 36°C or 29°C.

To quantify the reversion frequency, double mutant cells were spread on YE5S plates at the density of 2×10^4^ or 2×10^5^ cells per plate, incubated 4-5 days at 36°C. The number of non-ts revertant colonies was scored in three biological repeats for each double mutant strains. Two non-ts revertants were picked for each strain and the *ssp1* locus in these revertants is PCR amplified/Sanger sequenced to verify true reversion of *ssp1*^*MTD-AGGCA*^ to wild type.

We also did a fluctuation test for some double mutant strains to quantify the reversion frequency by growing a single overnight culture in YE5S broth for the strain to be tested, diluting with fresh YE5S broth to obtain 10^2^ yeast cells /ml. For each strain, the diluted suspension was divided into 48 of 100μL and incubated at 29°C. Then 40 replica cultures with 100μL were plated in their entirety onto YE5S agar plates and incubated at 36°C for 3∼5 days. For the rest 10 100μL replica cultures, the average number of cells per culture (N) was calculated using a blood counting chamber. Then counted the number of 36°C survival cells (reverted wt cells) per culture, and calculated the mutation rate with the p0 method or the MSS-maximum likelihood method (67).

### Identification of *yox1*^MTD^ and *lsk1*^MTD^ among *cnp1*^*H100M*^ suppressors and stability test for *yox1*^MTD^

Haploid *cnp1*^*H100M*^ cells derived from heterozygous *cnp1*^*H100M*^ diploid by tetrad dissection were spread on YE5S plates, incubated at 29°C for 5 days. Rare large colonies (∼1/10^4^) were isolated as spontaneous *cnp1*^*H100M*^ suppressors (FigS2A). Whole-genome sequencing was performed on isolated *cnp1*^*H100M*^ suppressors to identify the target gene. With the analysis process in “Whole-genome sequencing and datasets analysis” part, MTD events in *yox1* and *lsk1* gene were identified and verified by Sanger sequencing.

To verify the genetic stability of *yox1*^MTD^ alleles, *cnp1*^*H100M*^ suppressors were backcrossed with wild type, *yox1*^MTD^ were separated from *cnp1*^H100M^ mutation. *yox1*-GFP, *yox1*^*MTD*^-GFP strains were constructed by fusing a GFP tag in the endogenous *yox1* locus (FigS2B). MTD (20bp tandem duplication) in *yox1* disrupts the open reading frame and generates a premature stop codon (TAG) at 523nt loci, resulting in inactivation of GFP fluorescence, while the reversion of yox1^MTD^ would recover the GFP fluorescence. In the stability test, *yox1*-GFP and *yox1*^MTD^-GFP cells were grown continuously at 29°C by refreshing the culture with YE5S liquid media daily for up to 60 days. Every ten days, cell samples were taken and subjected to microscopical observation for GFP fluorescence. The percentage of progenies exhibiting the nuclear GFP signal was scored in three individual biological repeats. To verify *yox1*^MTD^-GFP reversion, *yox1* locus of @ single colonies derived from *yox1*^MTD^-GFP 40 day culture was amplified by PCR and subjected for Sanger sequencing

### Finding microhomology pairs on genome

A fast algorithm is implemented to find micro-homology pairs across the S. pombe’s genome sequence (or any given DNA sequence). First, the input sequence is scanned one-time for initial k-mer homology pairs with pre-set limitations. Here we arbitrarily set limitations to 1) the size of the homology should be no smaller than 4 bps and no greater than 12 bps, 2) the homology should not be a mononucleotide repeat, 3) space between two homologies in a pair should be greater than 3 bps, and 4) the INDEL size (the length of a homology plus the inter-space) should not exceed 100 bps. Then, the initial homology pairs are forth scanned for one run to merge adjacent homology pairs to longer pairs. The current implementation would only report the left-most pair of tandem repeats with micro-homology pairs on repeat junctions

### Annotating insertions and tandem repeats flanked by micro-homology pairs in natural isolates and in the reference genome

To identify MH-flanked tandem repeats in the reference genome we used the Tandem Repeat Finder(68) to generate an initial tandem repeat candidate list. All parameters were set to the default value except the INDEL penalty, which was set to 1000 to avoid reporting tandem repeats with non-uniform unit sizes. After removing candidates with the reported unit size smaller than 10nt, self-information smaller than 1.5 bits, and repeat number smaller than 2, remaining tandem repeats were verified by three steps: 1) finding if there were still internal repeats within the reported repeat unit, 2) finding if there were still repeat units on the left and right wings to the reported length, and 3) sliding the whole frame to the left-most base while the repeats’ consistency did not drop. Finally, we checked the junctions for the existence of a micro-homology of at least 2nt. If homology size is long enough (longer than 75% of the unit size and longer than unit size–4 bps), we considered it as repeat number plus 1 and start over for finding junction micro-homologies.

To identify MH-flanked insertions in natural isolates we used the indels .vcf file from Jeffares et al.(42) and used SnpEff (69) to predict the impact of each indel. We extracted the left and right flanking sequences from the reference genome to determine the presence of microhomology and to identify the repeat unit.

### Site-specific PCR amplification and ultra-deep next-generation sequencing

The fresh single colony was picked from the YE5S plate, inoculate 3ml YE5S liquid medium and incubated at 29°C overnight. The mini-culture was refreshed by 1:10 dilution in 20ml YE5S liquid medium and grown to mid-log phase. Genomic DNA was extracted using phenol-chloroform, used as the template for PCR amplification with high fidelity polymerase (RR006Q, Takara, Tokyo, Japan). Alternatively, a plasmid containing the *ssp1* coding sequence was constructed and amplified in E.coli, extracted, and digested with endonucleases to release the *ssp1* DNA fragments. Chemically synthesized ssp1 DNA fragments were produced by commercial service (Hzykang, Hangzhou, China). ssp1 DNA fragments from various sources described above were subjected to Illumina NGS following standard procedure at the coverage of ∼1×10^6^ (Bioacme, Wuhan, China).

The sequencing library of wild type diploid cells derived from a single colony was prepared as above and subjected to Illumina NGS by Frasergen in Wuhan, China.

For sequencing data analysis, trimmed FASTQ files are mapped to the reference sequences with BWA mem (with -Y flag on) and only primary alignments are kept. The program described in “Finding micro-homology pairs” is used here to find micro-homology pairs in the library reference. The left and right adjacent bases (here we arbitrarily chose 10 bps) to each of the two homologies in the micro-homology pairs are extracted as “signature sequences”. Then the alignment maps are scanned: for a clipped read, in those pairs that are possible to generate duplication/collapse at the clipping position, we test whether the clipped sequence matches with any pair’s “signature sequence”; for an INDEL possessing read, we test the opening and ending positions (and as well the inserted sequence for insertion reads).

### Simulations of mutation frequency at different reversion rates

Let *A*_*wt*_ be the wild-type allele, and *A*_*mut*_ be the mutant allele. Let *k*_*fwd*_ be the forward (*A*_*wt*_ to *A*_*mut*_) mutation rate, and *k*_*rev*_ be the reverse (*A*_*mut*_ to *A*_*wt*_) mutation rate. Let *p*_*t*_ be the frequency of *A*_*wt*_ and *q*_*t*_ the frequency of *A*_*mut*_ at time *t*. Then, if we assume that mutations are neutral, the *A*_*mut*_ genotype frequency (*q*) changes as *q*_(*t+1*)_ *= q*_*t*_ *+* (*k*_*fwd*_ × *p*_*t*_ *– k*_*rev*_ × *q*_*t*_), and *p*=(1-*q*). When *k*_*fwd*_ >= *k*_*rev*_ or, as is the case for subclonal MTDs, when *q* is small, the reverse mutation can mostly be ignored. However, when *k*_*rev*_ ≫ *k*_*fwd*_ or *q* == 1, as is the case for clonal fixed MTDs, *k*_*rev*_ has a large impact on dynamics. For simulations, the initial conditions were set to *p*=1,*q*=0, *k*_*fwd*_ =10^−7^, and *k*_*rev*_ was varied as is shown in the figure.

### Logistic regression to predict MTD frequency from local features

To predict the likelihood of a duplication event in each micro-homology pair (MHP), we used a logistic regression model (the function glm() from R) with 10-fold cross-validation. The data are highly imbalanced; MTDs were detected at fewer than 0.1% of MHPs. We therefore trained and tested the model using a balanced dataset consisting of all MHPs with an MTD, plus a randomly chosen subset MHPs with no MTD of the same size, so that half of MHPs had an MTD. We first trained a model using three features: MHlength, GC-content-of-the-MH-sequence, and inter-MH-distance, which has an AUC of 0.876. This is the “top 3 features” model, and all three of these features are predictive by visual inspection (e.g., Fig 2). To determine which additional features to add we continuously added features, and kept only those that increased the AUC over this base 3-feature model. The additional predictive features were: MHPlength (MHlen), nucleotides between two repeats (interMH), interMH (interGCcon), nucleosome occupancy (entire_nucle) and gene expression (entire_gene) of the entire MHP, and nucleotides to the closest MHR which has duplication event(ntclosestMHR).

To perform whole-genome predictions using the model trained on the balanced data, we used the model to score all 25 million MHPs in the genome, and either used the sum of predicted scores for all MHPs in a single gene, or selected the top 6234 MHPs, the same number of duplication events as observed experimentally, to be predicted duplication events.

**Figure S1.**
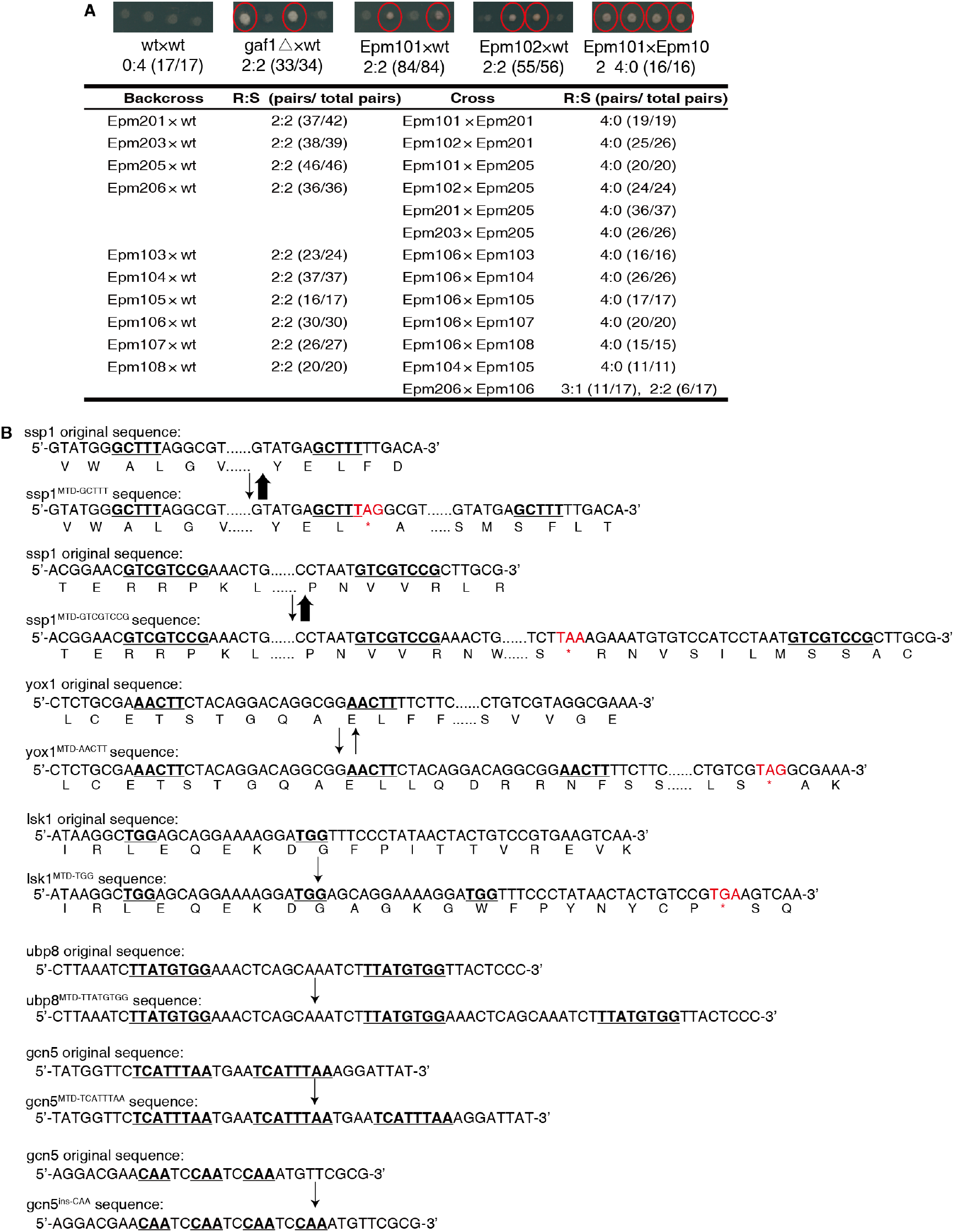
Identified highly reversible MTD mutations. **(A)**. Genetic linkage test for isolated reversible mutants in rapamycin plus caffeine screen. The ratio of resistant to sensitive progenies(R:S) is scored. The resistant progeny is labeled with red ellipses in the image panel above the table. And the statistic number in the brackets showed the pairs meeting the indicated R:S ratio/the total calculated pairs. **(B)**. Tandem duplication in multiple sites results in frame shift and pre-mature stop codon.

**Figure S2.**
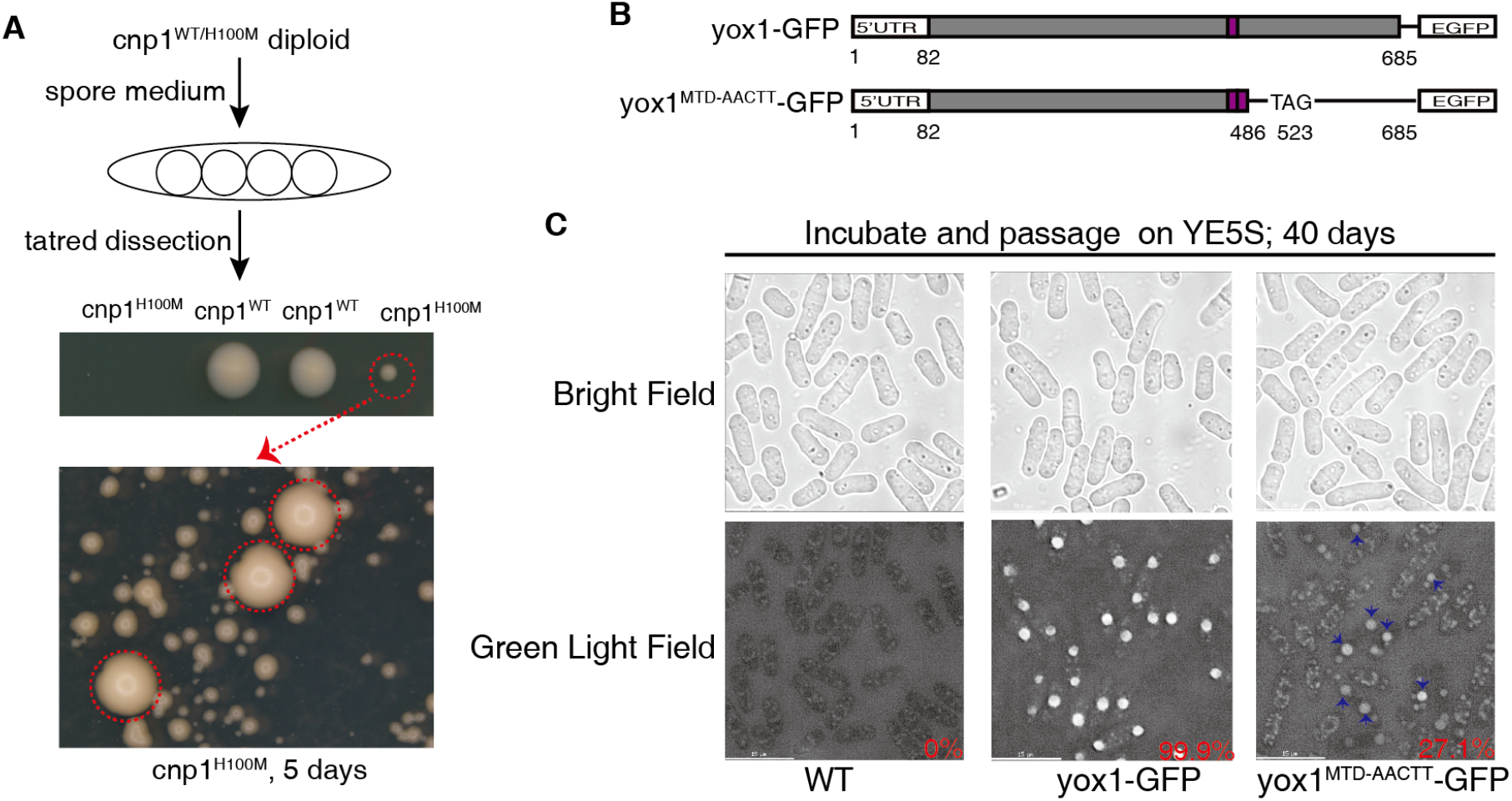
Identification of MTDs in *cnp1*^*H100M*^ suppressor screen. **(A)**. Process to isolate suppressors rescuing severe growth defect of *cnp1*^*H100M*^. Suppressors occurred after 5 days cultivation of cnp1^H100M^ mini-clones on YE5S plate, and marked with red dotted circle. **(B)**. Construction of yox1-GFP and yox1^MTD^-GFP strains. “TAG” is the premature stop codon. **(C)**. Genetic instability of yox1^MTD^ mutation is verified by fusing a GFP fluorescence marker. The blue arrows point recovered GFP signal, and the percentage marked with red shows the rate of cells with GFP signal.

**Figure S3.**
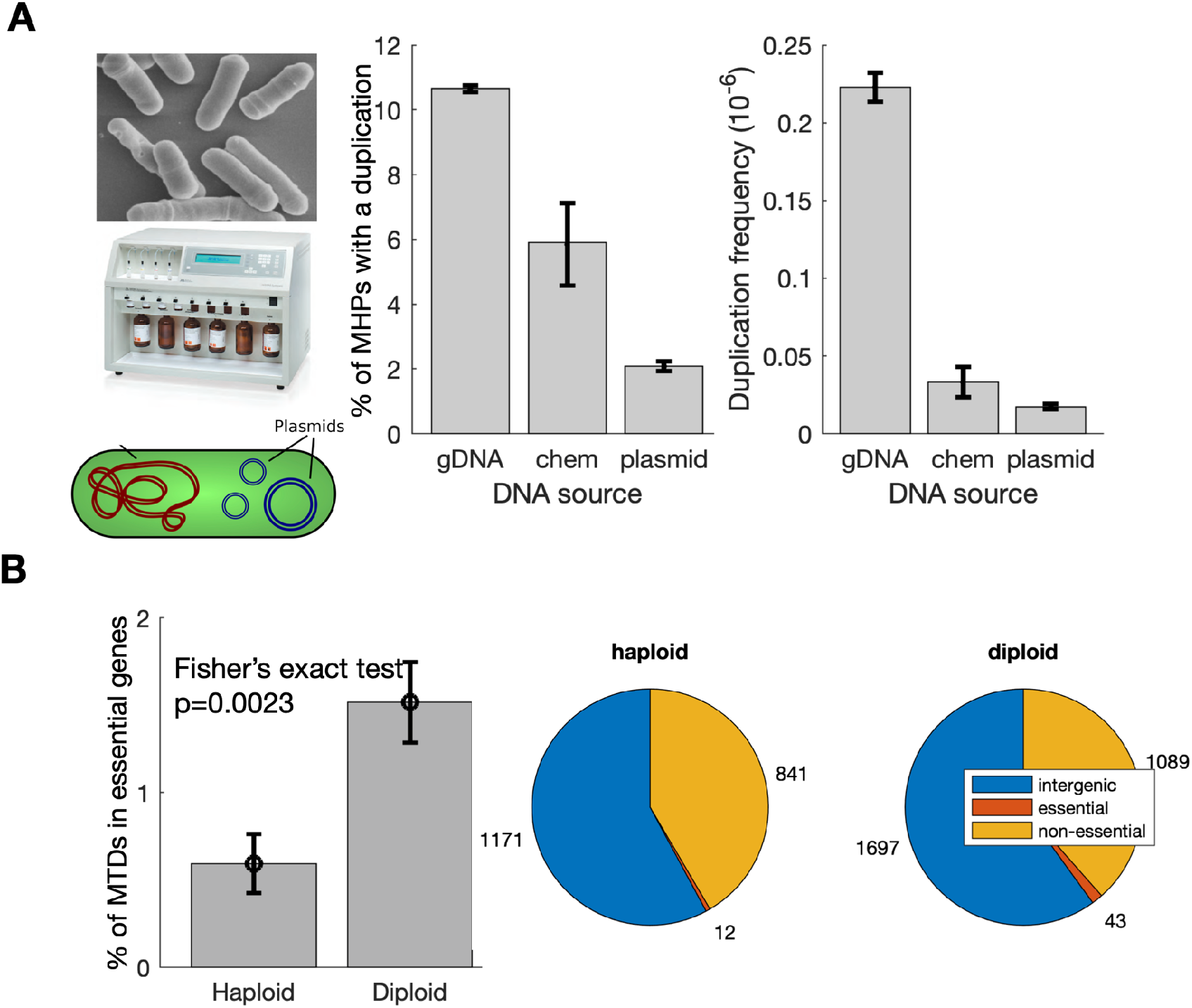
MTDs are more commonly observed in genomic DNA and subclonal MTDs in essential genes are more common in diploids in S. cerevisiae. **(A)** The *ssp1* gene was cloned into a plasmid in E. coli, and the gene was amplified by PCR from either *S. pombe* genomic DNA or miniprepped plasmid, or 200nt or 500nt chemically synthesized fragments, and all PCR amplicons were sequenced together to similar sequencing depths (10^5^-10^6^x coverage). Shown are the % of MHPs in *ssp1* in which a duplication was observed, as well as the measured duplication frequency (reads per 10^6^ coverage at that position). **(B)** Shown are the % of observed MTDs that are fully contained within essential genes in haploid or diploid mutation accumulation lines of budding yeast, as well as the distribution of MTDs throughout the genome. Reads from each haploid or diploid mutation accumulation line were mapped and analyzed independently, and the results merged.

**Figure S4.**
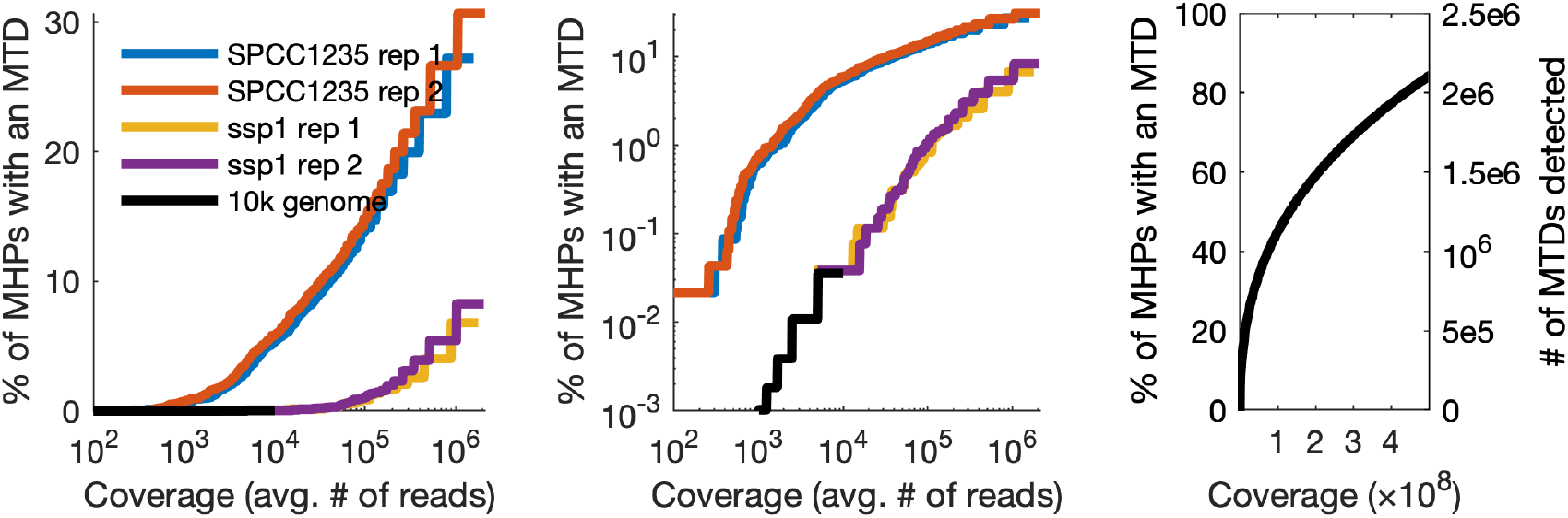
The measured (left & middle) and estimated (right) sequencing coverage required to observe all of the possible MTDs in the genome. Shown are the % of MHPs with an observed MTD in ultra-deep amplicon sequencing (single genes) and for 10k whole-genome sequencing (black) as a function of the sequencing coverage. SPCC1235 is a hot gene; the same coverage results in far more observed MTDs, while ssp1 is more representative of the genome as a whole. The far right shows simulated data where the 10k data + ssp1 line is extended out to 10^8^ coverage.

**Figure S5.**
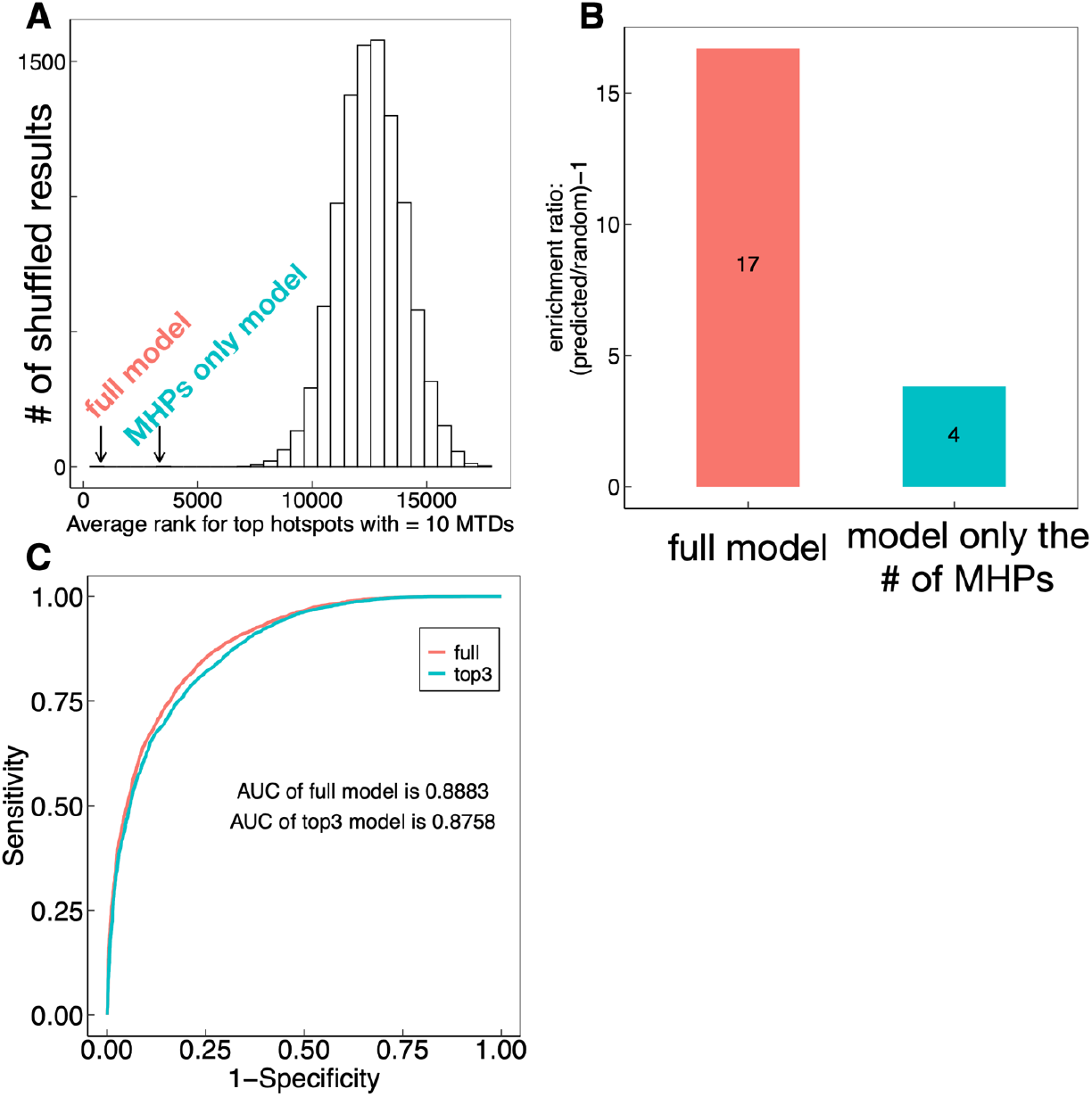
Characterization of the logistic regression model for predicting MTDs and hot-spots from *cis* MHP features. **(A,B)** Hotspots were defined as 1kb windows with more than 10 observed MTDs in the 10k whole-genome sequencing data. To determine if hotspots are solely due to MHP density, or are due to other sequence features incorporated into the model, we generate a random background distribution (histogram, white bars). The observed MTDs were shuffled across all MHPs in the genome, and the 1kb windows were ranked by the number of MTDs contained within each window (rank=1 has the most MTDs), and the average rank of the top windows was calculated. The classification model was then used to predict hotspots using all features, or only by counting MHPs. **(C)** Receiver Operating Characteristic (ROC) curve for models with all features, or with only GC content, inter-MH-distance, and MH length. the number of MHPs in each 1kb window. The full classification model outperforms the MHP count; hotspots are determined by more than just MHP density.

**Figure S6.**
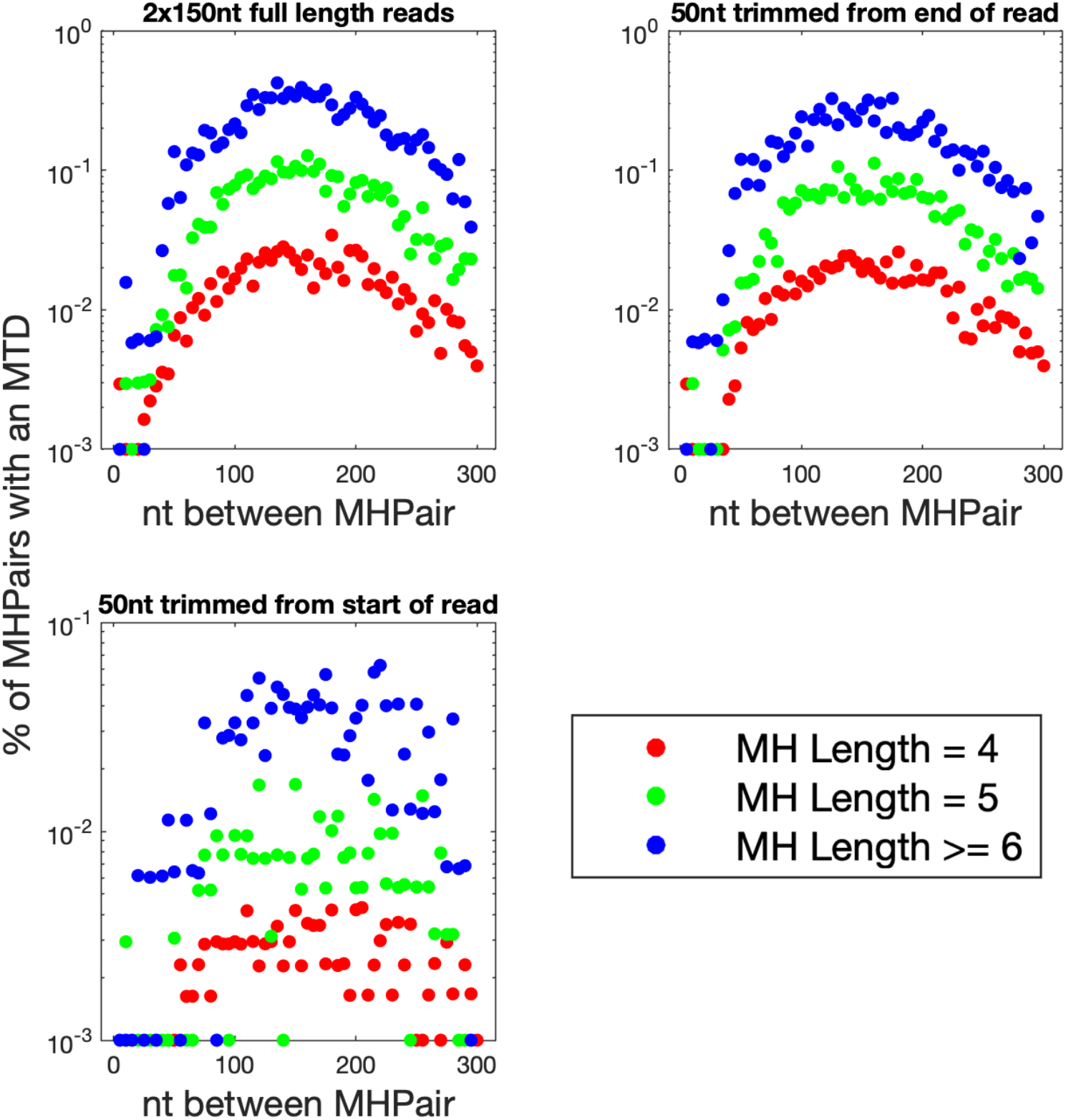
The relation between inter-MH spacing and MTD frequency is independent of read length. Trimmomatic(70) was used to remove either the first 50nt or the last 50nt from the end of each read, resulting in 2×100nt reads instead of 2×150nt reads; the peak at 150 remains unchanged. The higher noise when removing 50nt from the start is due to fewer identified MTDs, likely due to the higher error rate at the end of the read combined with the requirement for a perfect match to the MTD signature.

**Figure S7.**
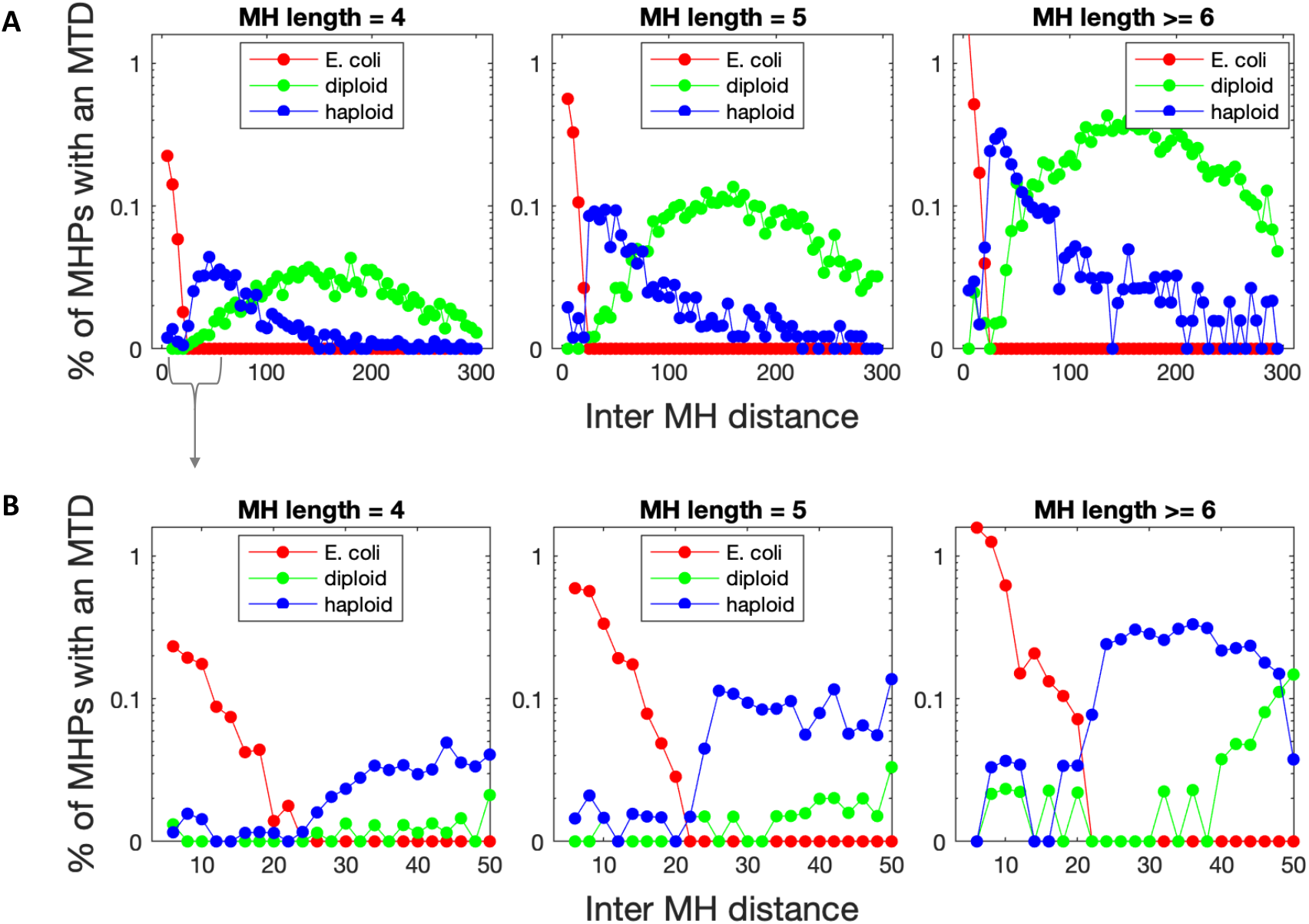
Characterization the relation between MH sequence length, inter-MH distance, and observed MTD frequency across different ultra-deep whole-genome sequencing datasets. **(A)** The relation between MTD frequency and inter-MH distance are shown for diploid *S. pombe* (green, this study), an isogenic haploid *S. pombe* (SRR7817502, 1700x coverage, blue), and E. coli (PRJNA329347, 14000x coverage, red). We note that the shorter haploid *S. pombe* (blue) inter-MH distance distribution is more similar to the insert lengths found in genetic screens, all of which were done in haploid strains. **(B)** Same data as in (A), but only inter-MH distances 3-50nt are shown.

**Figure S8.**
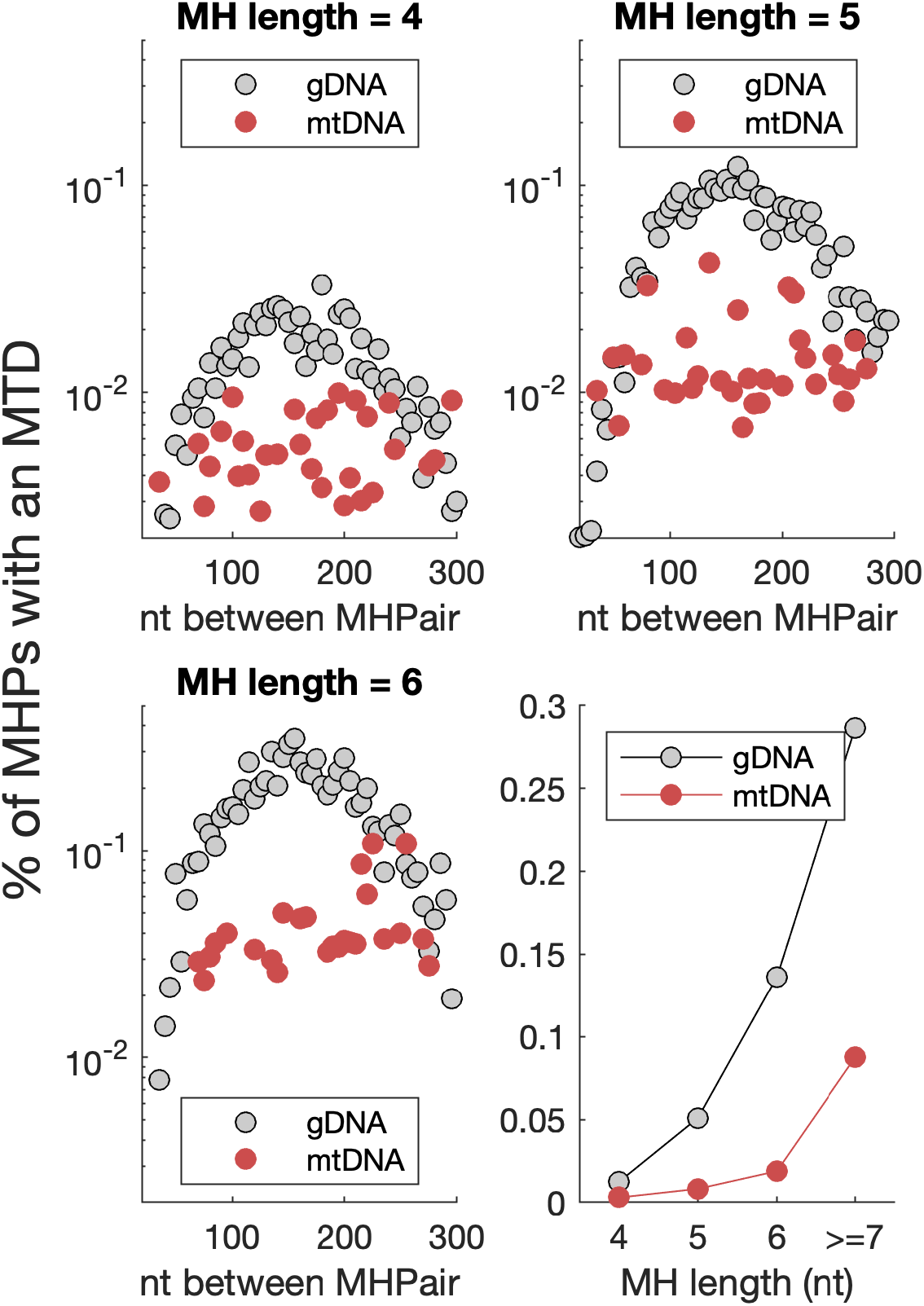
MTDs are less common in the mitochondria, and do not exhibit a peak at 150nt. MTDs in the mitochondrial DNA were downsampled so that the median sequencing coverage was identical to that of the gDNA. Downsampling was repeated 5000 times to increase the statistical power.

**Figure S9.**
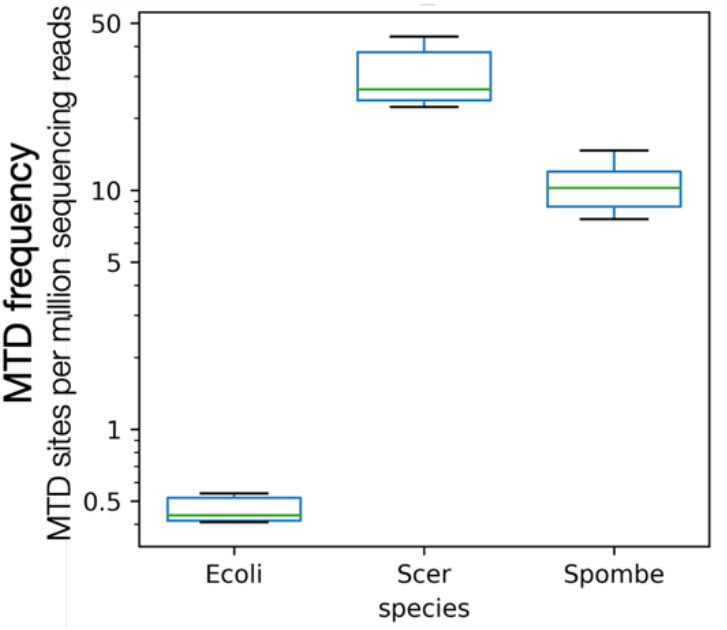
MTDs are 20x less common in the E. coli genome. Genomic DNA sequencing libraries of *E. coli, S. cerevisiae*, and *S. pombe* were prepared identically and sequenced together in a single sequencing lane, with multiple biological and technical replicates. For each replicate, the total number of MHPs with an MTD was divided by the sequencing depth. As the density of MHPs in the three genomes are roughly similar, normalizing by the number MHPs does not affect the result.

**Figure S10.**
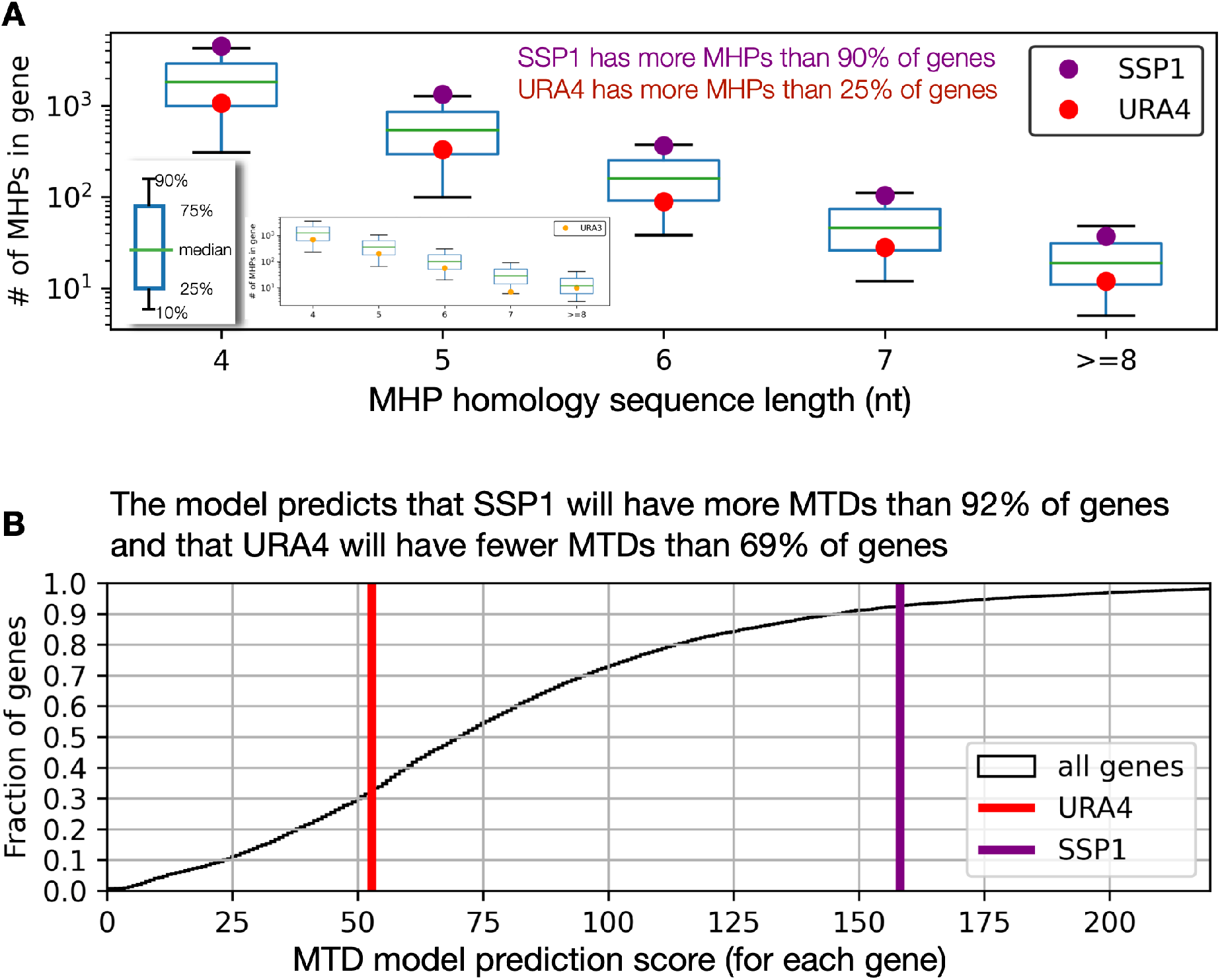
SSP1 is enriched for MHPs, and URA4 is depleted for MHPs. URA4, a common target of mutation screens, has relatively few MHPs. (A) The total number of MHPs in all genes in the genome (boxplot) with URA4 and SSP1 highlighted. Insert shows data from *S. cerevisiae* and URA3. (B) While the total number of MHPs is the size of the mutational target, URA4 is still depleted in MHPs even after normalizing by gene length. (C) The sequence-feature based model predicts that MTDs in URA4 will occur less often than in 70% of other genes.

**Figure S11.**
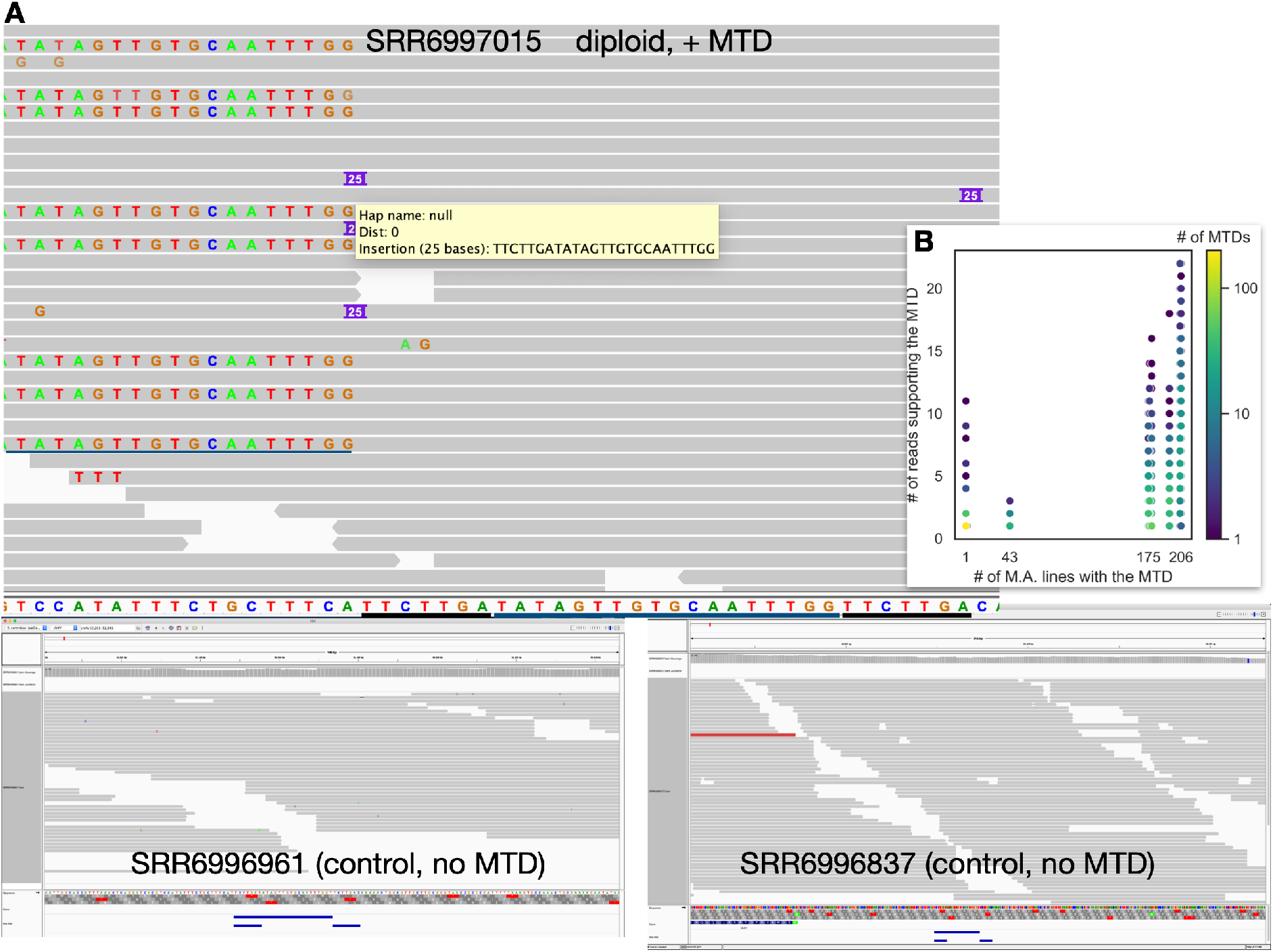
An example heterozygous MTD from an M.A. line. **(A)** One M.A. line that is heterozygous for an MTD in chrIV; this was called in (32), but not identified as an MTD. Reads from two other strains that do not have this tandem duplication are shown as controls. The MH-sequences are underlined in black, and the duplicated region in blue. **(B)** The distribution of the number of reads supporting each MTD (y-axis), for MTDs found in different numbers of the M.A. lines (x-axis). Color shows the number of different MTDs at each point.

**Figure S12.**
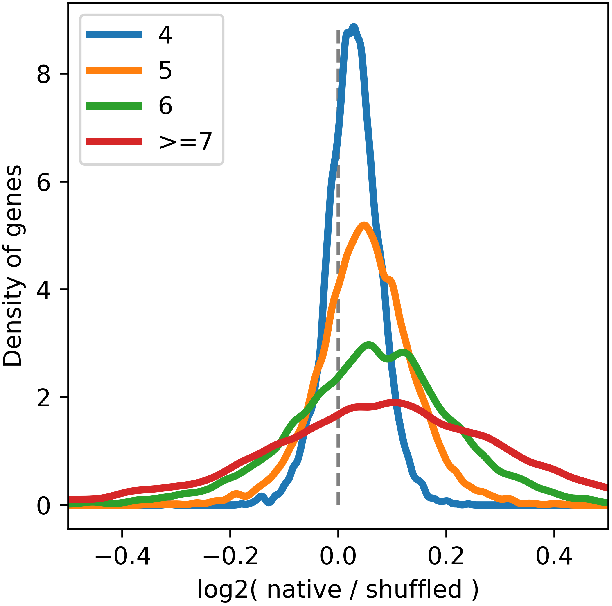
MHPs are more common in native genes that is expected by chance. For each protein coding gene in *S. pombe*, the number of MHPs with each MH sequence length (native) was compared to the average number of MHPs calculated across ten randomly generated synonymous versions of the protein, all of which maintain the same amino acid sequence, codon usage, and GC content. There was no difference between essential and non-essential genes.

## Supplementary Tables

**Supplementary Table S1**. Yeast strains used in this study.

**Supplementary Table S2**. All insertions and deletions identified in wild *S. pombe* isolates.

**Supplementary Table S3**. All indels >= 10bp identified in wild *S. pombe* isolates.

**Supplementary Table S4**. The number of MHPs and MTDs in each gene identified in the 10k ultra-deep sequencing data.

**Supplementary Table S5**. Features found to be predictive of MTD frequency.

**Supplementary Table S6**. All single gene deletion mutants screened in the reversion assay.

**Supplementary Table S7**. All MTDs identified in all genetic screens.

## Notes

### Competing Interest Statement

The authors have declared no competing interest.

### Summary of Updates

new data and analysis for E. coli lots of new citations to place the paper in context an extended discussion

## References and Notes

1. O. J. Rando, K.J. Verstrepen, Timescales of genetic and epigenetic inheritance. Cell 128, 655–668 (2007).

2. D. I. Andersson, B. R. Levin, The biological cost of antibiotic resistance. Curr. Opin. Microbiol. 2, 489– 493 (1999).

3. R. E. Lenski, Bacterial evolution and the cost of antibiotic resistance. Int. Microbiol. 1, 265–270 (1998).

4. R. Gemayel, M. D. Vinces, M. Legendre, K. J. Verstrepen, Variable tandem repeats accelerate evolution of coding and regulatory sequences. Annu. Rev. Genet. 44, 445–477 (2010).

5. J. E. Haber, E. J. Louis, Minisatellite Origins in Yeast and Humans. Genomics 48, 132–135 (1998).

6. K. J. Verstrepen, A. Jansen, F. Lewitter, G. R. Fink, Intragenic tandem repeats generate functional variability. Nat Genet 37, 986–990 (2005).

7. E. R. Moxon, P. B. Rainey, M. A. Nowak, R. E. Lenski, Adaptive evolution of highly mutable loci in pathogenic bacteria. Curr Biol 4, 24–33 (1994).

8. R. Moxon, C. Bayliss, D. Hood, Bacterial contingency loci: the role of simple sequence DNA repeats in bacterial adaptation. Annu Rev Genet 40, 307–333 (2006).

9. S. Calo, et al., Antifungal drug resistance evoked via RNAi-dependent epimutations. Nature 513, 555–558 (2014).

10. S. M. Shaffer, et al., Rare cell variability and drug-induced reprogramming as a mode of cancer drug resistance. Nature 546, 431–435 (2017).

11. S. V. Sharma, et al., A Chromatin-Mediated Reversible Drug-Tolerant State in Cancer Cell Subpopulations. Cell 141, 69–80 (2010).

12. R. Dhar, A. M. Missarova, B. Lehner, L. B. Carey, Single cell functional genomics reveals the importance of mitochondria in cell-to-cell phenotypic variation. Elife 8 (2019).

13. S. F. Levy, N. Ziv, M. L. Siegal, Bet hedging in yeast by heterogeneous, age-correlated expression of a stress protectant. PLoS Biol. 10, e1001325 (2012).

14. H. Nicoloff, K. Hjort, B. R. Levin, D. I. Andersson, The high prevalence of antibiotic heteroresistance in pathogenic bacteria is mainly caused by gene amplification. Nat Microbiol 4, 504–514 (2019).

15. R. T. Todd, A. Selmecki, Expandable and reversible copy number amplification drives rapid adaptation to antifungal drugs. Elife 9 (2020).

16. T. D. Tlsty, A. M. Albertini, J. H. Miller, Gene amplification in the lac region of E. coli. Cell 37, 217–224 (1984).

17. T. Huang, J. L. Campbell, Amplification of a Circular Episome Carrying an Inverted Repeat of the DFR1 Locus and Adjacent Autonomously Replicating Sequence Element of Saccharomyces cerevisiae. J. Biol. Chem. 270, 9607–9614 (1995).

18. P. J. Hastings, H. J. Bull, J. R. Klump, S. M. Rosenberg, Adaptive Amplification. Cell 103, 723–731 (2000).

19. D. L. Hartl, E. W. Jones, Genetics: principles and analysis, 4th ed (Jones and Bartlett Publishers, 1998).

20. R. Lande, Risk of population extinction from fixation of deleterious and reverse mutations. Genetica 102–103, 21–27 (1998).

21. T. Maruyama, M. Kimura, A NOTE ON THE SPEED OF GENE FREQUENCY CHANGES IN REVERSE DIRECTIONS IN A FINITE POPULATION. Evolution 28, 161–163 (1974).

22. L. B. Carey, RNA polymerase errors cause splicing defects and can be regulated by differential expression of RNA polymerase subunits. Elife 4 (2015).

23. R. Weisman, M. Choder, Y. Koltin, Rapamycin specifically interferes with the developmental response of fission yeast to starvation. J. Bacteriol. 179, 6325–6334 (1997).

24. D. Laor, A. Cohen, M. Kupiec, R. Weisman, TORC1 Regulates Developmental Responses to Nitrogen Stress via Regulation of the GATA Transcription Factor Gaf1. mBio 6, e00959 (2015).

25. E. Davie, G. M. A. Forte, J. Petersen, Nitrogen regulates AMPK to control TORC1 signaling. Curr. Biol. 25, 445–454 (2015).

26. A. R. J. Lawson, et al., RAF gene fusion breakpoints in pediatric brain tumors are characterized by significant enrichment of sequence microhomology. Genome Res. 21, 505–514 (2011).

27. L. E. L. M. Vissers, et al., Rare pathogenic microdeletions and tandem duplications are microhomology-mediated and stimulated by local genomic architecture. Hum. Mol. Genet. 18, 3579–3593 (2009).

28. N. A. Willis, et al., Mechanism of tandem duplication formation in BRCA1-mutant cells. Nature 551, 590– 595 (2017).

29. E. Harrison, V. Koufopanou, A. Burt, R. C. MacLean, The cost of copy number in a selfish genetic element: the 2-μm plasmid of Saccharomyces cerevisiae. J Evol Biol 25, 2348–2356 (2012).

30. E. M. Torres, et al., Effects of aneuploidy on cellular physiology and cell division in haploid yeast. Science 317, 916–924 (2007).

31. S. R. Head, et al., Library construction for next-generation sequencing: overviews and challenges. BioTechniques 56, 61–64, 66, 68, passim (2014).

32. N. P. Sharp, L. Sandell, C. G. James, S. P. Otto, The genome-wide rate and spectrum of spontaneous mutations differ between haploid and diploid yeast. Proc Natl Acad Sci USA 115, E5046–E5055 (2018).

33. A. Baryshnikova, et al., Quantitative analysis of fitness and genetic interactions in yeast on a genome scale. Nat. Methods 7, 1017–1024 (2010).

34. A. Slack, P. C. Thornton, D. B. Magner, S. M. Rosenberg, P. J. Hastings, On the mechanism of gene amplification induced under stress in Escherichia coli. PLoS Genet. 2, e48 (2006).

35. S. Gangloff, et al., Quiescence unveils a novel mutational force in fission yeast. eLife 6, e27469 (2017).

36. X. Xu, L. Wang, M. Yanagida, Whole-Genome Sequencing of Suppressor DNA Mixtures Identifies Pathways That Compensate for Chromosome Segregation Defects in Schizosaccharomyces pombe. G3 8, 1031–1038 (2018).

37. I. Iraqui, et al., Recovery of arrested replication forks by homologous recombination is error-prone. PLoS Genet. 8, e1002976 (2012).

38. S. T. Lovett, Encoded errors: mutations and rearrangements mediated by misalignment at repetitive DNA sequences. Mol Microbiol 52, 1243–1253 (2004).

39. A. Teixeira-Silva, et al., The end-joining factor Ku acts in the end-resection of double strand break-free arrested replication forks. Nat Commun 8, 1982 (2017).

40. A. A. Alcasabas, et al., Mrc1 transduces signals of DNA replication stress to activate Rad53. Nat. Cell Biol. 3, 958–965 (2001).

41. K. Myung, R. D. Kolodner, Suppression of genome instability by redundant S-phase checkpoint pathways in Saccharomyces cerevisiae. Proc. Natl. Acad. Sci. U.S.A. 99, 4500–4507 (2002).

42. D. C. Jeffares, et al., The genomic and phenotypic diversity of Schizosaccharomyces pombe. Nat Genet 47, 235–241 (2015).

43. C. Duan, et al., Reduced intrinsic DNA curvature leads to increased mutation rate. Genome Biol. 19, 132 (2018).

44. F. Supek, B. Lehner, Scales and mechanisms of somatic mutation rate variation across the human genome. DNA Repair (Amst) 81, 102647 (2019).

45. D. Ottaviani, M. LeCain, D. Sheer, The role of microhomology in genomic structural variation. Trends Genet. 30, 85–94 (2014).

46. M. McVey, S. E. Lee, MMEJ repair of double-strand breaks (director’s cut): deleted sequences and alternative endings. Trends in Genetics 24, 529–538 (2008).

47. E. Darmon, D. R. F. Leach, Bacterial genome instability. Microbiol Mol Biol Rev 78, 1–39 (2014).

48. J. N. Vaughn, J. L. Bennetzen, Natural insertions in rice commonly form tandem duplications indicative of patch-mediated double-strand break induction and repair. Proc Natl Acad Sci U S A 111, 6684–6689 (2014).

49. M. Bzymek, S. T. Lovett, Instability of repetitive DNA sequences: The role of replication in multiple mechanisms. Proceedings of the National Academy of Sciences 98, 8319–8325 (2001).

50. P. J. Hastings, G. Ira, J. R. Lupski, A Microhomology-Mediated Break-Induced Replication Model for the Origin of Human Copy Number Variation. PLOS Genetics 5, e1000327 (2009).

51. D. X. Tishkoff, N. Filosi, G. M. Gaida, R. D. Kolodner, A novel mutation avoidance mechanism dependent on S. cerevisiae RAD27 is distinct from DNA mismatch repair. Cell 88, 253–263 (1997).

52. A. M. Albertini, M. Hofer, M. P. Calos, J. H. Miller, On the formation of spontaneous deletions: the importance of short sequence homologies in the generation of large deletions. Cell 29, 319–328 (1982).

53. T. Edlund, S. Normark, Recombination between short DNA homologies causes tandem duplication. Nature 292, 269–271 (1981).

54. H. Xia, W. Zhao, Y. Shi, X.-R. Wang, B. Wang, Microhomologies Are Associated with Tandem Duplications and Structural Variation in Plant Mitochondrial Genomes. Genome Biol Evol 12, 1965–1974 (2020).

55. S. K. Whoriskey, V. H. Nghiem, P. M. Leong, J. M. Masson, J. H. Miller, Genetic rearrangements and gene amplification in Escherichia coli: DNA sequences at the junctures of amplified gene fusions. Genes & Development 1, 227–237 (1987).

56. R. J. Kaufman, P. C. Brown, R. T. Schimke, Amplified dihydrofolate reductase genes in unstably methotrexate-resistant cells are associated with double minute chromosomes. Proceedings of the National Academy of Sciences 76, 5669–5673 (1979).

57. C. Payen, et al., The dynamics of diverse segmental amplifications in populations of Saccharomyces cerevisiae adapting to strong selection. G3 (Bethesda) 4, 399–409 (2014).

58. S. Mitsuhashi, et al., Tandem-genotypes: robust detection of tandem repeat expansions from long DNA reads. Genome Biol 20, 58 (2019).

59. D.-U. Kim, et al., Analysis of a genome-wide set of gene deletions in the fission yeast Schizosaccharomyces pombe. Nat. Biotechnol. 28, 617–623 (2010).

60. M. D. Krawchuk, W. P. Wahls, High-efficiency gene targeting in Schizosaccharomyces pombe using a modular, PCR-based approach with long tracts of flanking homology. Yeast 15, 1419–1427 (1999).

61. K. Ekwall, G. Thon, Spore Analysis and Tetrad Dissection of Schizosaccharomyces pombe. Cold Spring Harb Protoc 2017, pdb.prot091710 (2017).

62. W. Escorcia, S. L. Forsburg, Tetrad Dissection in Fission Yeast. Methods Mol Biol 1721, 179–187 (2018).

63. H. Li, Aligning sequence reads, clone sequences and assembly contigs with BWA-MEM. arXiv:1303.3997 [q-bio] (2013) (December 15, 2020).

64. H. Li, et al., The Sequence Alignment/Map format and SAMtools. Bioinformatics 25, 2078–2079 (2009).

65. A. McKenna, et al., The Genome Analysis Toolkit: a MapReduce framework for analyzing next-generation DNA sequencing data. Genome Res 20, 1297–1303 (2010).

66. W. McLaren, et al., The Ensembl Variant Effect Predictor. Genome Biol 17, 122 (2016).

67. G. I. Lang, Measuring Mutation Rates Using the Luria-Delbrück Fluctuation Assay. Methods Mol. Biol. 1672, 21–31 (2018).

68. G. Benson, Tandem repeats finder: a program to analyze DNA sequences. Nucleic Acids Res. 27, 573–580 (1999).

69. P. Cingolani, et al., A program for annotating and predicting the effects of single nucleotide polymorphisms, SnpEff: SNPs in the genome of Drosophila melanogaster strain w1118; iso-2; iso-3. Fly (Austin) 6, 80–92 (2012).

70. A. M. Bolger, M. Lohse,B. Usadel, Trimmomatic: a flexible trimmer for Illumina sequence data. Bioinformatics 30, 2114–2120 (2014).

